# Cell cycle-regulated transcriptional pausing of *Drosophila* replication-dependent histone genes

**DOI:** 10.1101/2024.12.16.628706

**Authors:** James P. Kemp, Mark S. Geisler, Mia Hoover, Chun-Yi Cho, Patrick H. O’Farrell, William F. Marzluff, Robert J. Duronio

## Abstract

Coordinated expression of replication-dependent (RD) histones genes occurs within the Histone Locus Body (HLB) during S phase, but the molecular steps in transcription that are cell cycle regulated are unknown. We report that *Drosophila* RNA Pol II promotes HLB formation and is enriched in the HLB outside of S phase, including G1-arrested cells that do not transcribe RD histone genes. In contrast, the transcription elongation factor Spt6 is enriched in HLBs only during S phase. Proliferating cells in the wing and eye primordium express full-length histone mRNAs during S phase but express only short nascent transcripts in cells in G1 or G2 consistent with these transcripts being paused and then terminated. Full-length transcripts are produced when Cyclin E/Cdk2 is activated as cells enter S phase. Thus, activation of transcription elongation by Cyclin E/Cdk2 and not recruitment of RNA pol II to the HLB is the critical step that links histone gene expression to cell cycle progression in *Drosophila*.

## Introduction

To properly replicate the genome during S phase of the cell cycle, eukaryotic cells must continuously supply sufficient histone proteins to package newly replicated DNA into chromatin. The bulk of these histone proteins (i.e. the 4 nucleosomal histones plus linker histone H1) are derived from multi-copy replication-dependent (RD) histone genes that lack introns and are clustered in metazoan genomes (Marzluff et al., 2008). These genes produce the only cellular mRNAs that are not polyadenylated but instead end in a conserved stemloop that is necessary for their coordinate regulation during the cell cycle (Marzluff and Koreski, 2017). RD-histone genes are present in a nuclear body called the Histone Locus Body (HLB), a biomolecular condensate which concentrates factors necessary for the synthesis of these unique mRNAs (Duronio and Marzluff, 2017; Geisler et al., 2023). The core factors and essential protein-protein interactions present in HLBs are conserved between *Drosophila* and mammals (Kemp et al., 2021; Skrajna et al., 2018; Yang et al., 2014). HLB assembly requires a large, mostly intrinsically disordered protein (NPAT in humans and Mxc in *Drosophila*) that oligomerizes and promotes liquid-liquid phase separation and is necessary for subsequent recruitment of other HLB components (Hur et al., 2020; Terzo et al., 2015; White et al., 2011). HLB assembly and histone gene expression do not occur in the absence of Mxc/NPAT (White et al., 2011; Ye et al., 2003). Mutations in the histone locus that prevent HLB formation abolish expression of all histone genes in the cluster (Chaubal et al., 2023; Koreski et al., 2020; Salzler et al., 2013). Furthermore, efficient RD histone pre-mRNA processing requires concentration of key factors, U7 snRNP and FLASH, in the HLB (Tatomer et al., 2016). Although the HLB is critical for RD histone gene expression, the mechanistic relationship between HLB formation and RD histone mRNA biosynthesis is not understood, nor are all the specific steps in histone mRNA biosynthesis that are regulated during the cell cycle.

*Drosophila melanogaster* RD histone genes are present in a ∼600 kB array containing ∼110 copies of each gene, representing about 0.3% of the genome (Bongartz and Schloissnig, 2019; Crain et al., 2024; McKay et al., 2015; Shukla et al., 2024).

Consequently, *Drosophila* HLBs are large and easily visualized within cells. RD histone gene transcription and pre-mRNA processing both occur within the HLB (Kemp et al., 2021). Because histone mRNA is rapidly exported to the cytoplasm after synthesis (Adesnik and Darnell, 1972), RD histone RNA visualized in the HLB by in situ hybridization represents nascent transcripts on histone genes. Consequently, the detection of nascent transcripts in HLBs allows one to readily visualize dynamic histone gene transcription in individual cells throughout the cell cycle (Guglielmi et al., 2013; Lanzotti et al., 2002; Lanzotti et al., 2004; Tatomer et al., 2016). Moreover, determining when and how mRNA biosynthetic factors are recruited to the HLB provides a powerful means to examine how biomolecular condensates contribute to gene expression as well as which steps are cell cycle regulated (Tatomer et al., 2014; Tatomer et al., 2016).

Once assembled, HLBs persist throughout interphase of the cell cycle, disassembling only during mitosis, and thus the assembly and disassembly of HLBs are not the primary means of restricting RD histone mRNA expression to S phase (Armstrong et al., 2023; Hur et al., 2020; Liu et al., 2006; Terzo et al., 2015; White et al., 2007). Some HLB factors (e.g. Mxc/NPAT, and the pre-mRNA processing factors FLASH and U7 snRNP) reside constitutively in the HLB (Liu et al., 2006; White et al., 2011), whereas others are recruited to the HLB only when histone mRNAs are expressed (e.g. the cleavage complex that assembles on the active U7 snRNP to cleave histone pre-mRNA) (Tatomer et al., 2014).

RD histone gene transcription plays a role in HLB formation and structure as it is necessary for *Drosophila* HLBs to attain their full size, but exactly what that entails mechanistically is unclear (Hur et al., 2020; Salzler et al., 2013). Transcription of RD histone genes requires both transcription factors that are specific for a type of histone gene (e.g. histone H1, H4, H2B) (Dailey et al., 1988; Fletcher et al., 1987; Gallinari et al., 1989; Lee et al., 2010; Mitra et al., 2003; Nirala et al., 2021) and the core RNA pol II machinery which is present in the HLB (Guglielmi et al., 2013). At individual genes, RNA pol II can exist in different states including bound to promoters but not transcribing, paused after initiating transcription, and actively elongating to synthesize mRNA (Adelman and Lis, 2012; Core et al., 2012). In addition, some RNA pol II localized to the HLB may not be bound to histone genes. A transition between any (or all) of these states could provide a key, cell cycle-regulated step that controls RD histone gene transcription. Mxc/NPAT localizes only to the promoters of RD histone genes in both human cells (Kaya-Okur et al., 2019) and *Drosophila* (Hodkinson et al., 2023), and phosphorylation of Mxc/NPAT by Cyclin E/Cdk2 is required for activation of histone mRNA synthesis (Armstrong et al., 2023; Lanzotti et al., 2004; Ma et al., 2000; Miele et al., 2005; Wei et al., 2003; White et al., 2007; Zhao et al., 2000). Accordingly, phosphorylation of Mxc/NPAT or other HLB components could regulate the recruitment of RNA pol II or other steps in transcription and may independently regulate activity of pre-mRNA processing factors. We don’t know which step(s) in RD histone mRNA synthesis are activated by phosphorylation of HLB components, but current evidence supports a role for Cyclin E/Cdk2 in both transcription and processing of RD histone mRNA (Armstrong et al., 2023; Hur et al., 2020; Tatomer et al., 2014).

Here we use live and fixed imaging of *Drosophila* tissues to explore the cell cycle regulation and dynamics of RNA polymerase II accumulation in the HLB at different stages of development. We find that RNA pol II accumulation in the HLB occurs outside of S phase and thus does not provide a means of coupling RD histone gene expression with the cell cycle. Instead, we provide evidence that release of paused RNA polymerase II into active elongation provides a major means of cell cycle control of expression of RD histone genes in *Drosophila*.

## Results

### RNA pol II and Spt6 have different dynamics of accumulation in the HLB during early *Drosophila* embryogenesis

To better understand the relationship between HLBs and RD histone mRNA synthesis, we performed a series of imaging experiments to compare the recruitment of RNA pol II components to the HLB and the dynamics of RD histone gene transcription. We first describe the features of RNA pol II recruitment to HLBs in several tissues during *Drosophila* development, followed by an analysis of the dynamics of nascent RD histone RNA within HLBs in the same tissues.

We began our analysis in embryos, which have a well characterized cell cycle program (Fig. 1A) (reviewed in (Swanhart et al., 2005; Yuan et al., 2016)). The first 13 nuclear division cycles of *Drosophila* embryogenesis are supported by maternally derived factors deposited into the egg, including histone proteins and mRNAs. These cycles occur synchronously in a syncytium and lack gap phases, progressing from S phase to mitosis and directly back into S phase. The first gap phase appears after S phase of the 14^th^ cycle, and during this G2 cellularization is completed and gastrulation begins. Patterned activation of Cdk1 coordinates entry into mitosis 14 with the events of gastrulation. Subsequent cell cycles 15 and 16 in the epidermis are also regulated at the G2-M transition and lack G1 phase, as S-phase begins immediately after mitosis 14 and mitosis 15. These post-blastoderm cell cycles differ from cell cycles later in development in that Cyclin E/Cdk2 remains active throughout interphase (including G2) until late in cycle 16 when it is down regulated (de Nooij et al., 1996; Lane et al., 1996). As a result of Cyclin E/Cdk2 downregulation during interphase of cycle 16, G1 phase first appears at cycle 17 in epidermal cells, and these cells remain G1-arrested until the embryo hatches into a first instar larva. Other embryonic lineages (e.g. neuroblasts) continue proliferating until late embryonic stages via G2-regulated cell cycles that lack a G1 phase.

**Figure 1.**
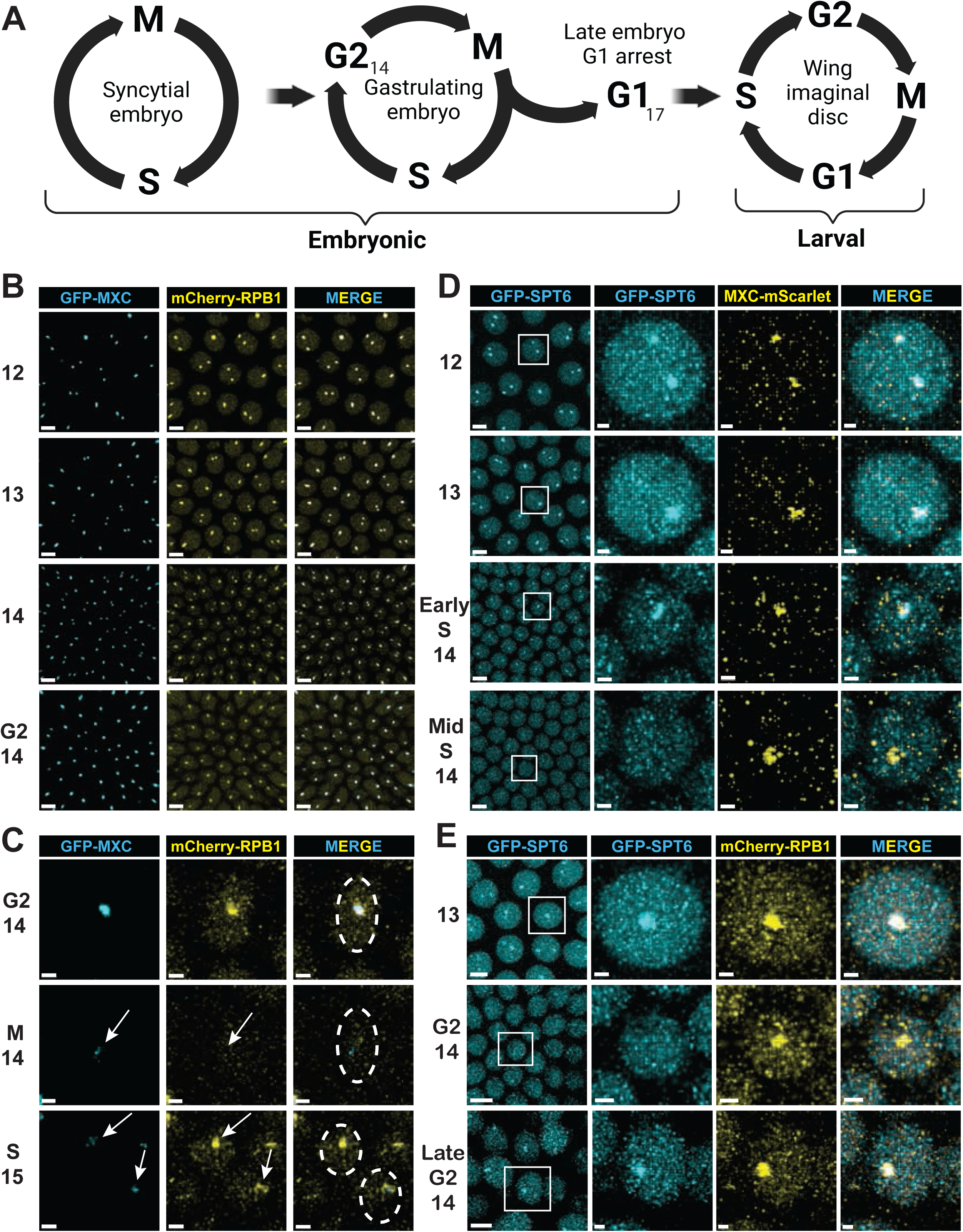
RNA pol II and Spt6 are recruited to the HLB with different dynamics. **A)** Diagram of the embryonic cell cycle program from rapid syncytial cycles to G2-regulated cycles in cycle 14 (G2_14_) to G1 arrest in epidermal cells at cycle 17 (G1_17_), followed by transition to canonical 4 phase cells cycles in the larval imaginal discs. Made with BioRender. **B)** Single micrographs from live imaging of syncytial nuclear cycles 12-14 in an embryo from a dual endogenously tagged strain containing both GFP-Mxc and mCherry-Rpb1. (Taken from supplemental videos 1A-C). Note the high level of RNA pol II within every HLB and that some cells have one HLB and some have 2 HLBs depending on whether the homologs have paired. Scale bars are 5 μM. **C)** Single micrographs from live imaging of a GFP-Mxc; mCherry-Rpb1 embryo as it begins gastrulation during G2 of cycle 14. (Taken from supplemental videos 2A-C). Arrows indicate disassembly of the HLB in mitosis 14 and rapid reformation in S phase of cycle 15. Dashed circle outlines the nucleus of the mother cell and the resulting daughter cells after mitosis. Scale bars are 2 μM. **D)** Single micrographs from live imaging of syncytial nuclear cycles 12-14 in an embryo from a dual endogenously tagged strain containing both GFP-Spt6 and Mxc-mScarlet. (Taken from supplemental videos 3A-C). Rows 3 and 4 show early-and mid-S phase of cycle 14, respectively. Scale bars are 5 and 1 μM in micrograph and magnified region of interest, respectively. **E)** Single micrographs from live imaging of cycles 13-14 in an embryo from a dual endogenously tagged strain containing GFP-Spt6 and mCherry-Rpb1. (Taken from supplemental videos 4-5A-C). Note cells in late G2 of cycle 14 have Spt6 in their HLBs. Scale bars are 5 and 1 μM in micrograph and region of interest, respectively.

The HLB first forms during syncytial nuclear cycle 11 (White et al., 2007) which is when zygotic RD histone mRNA synthesis is initiated (Edgar and Schubiger, 1986). We used live imagining of embryos that express both GFP-tagged Mxc and mCherry-tagged Rpb1, the large subunit of RNA pol II, from the endogenous genes to visualize both the HLB and RNA pol II during cycles 12-15. Previously, we and others found that throughout interphase of the syncytial cycles RNA pol II is enriched in nuclear foci, the largest of which coincide with Mxc and thus are HLBs (Fig. 1B, Supplemental Video 1A-C) (Blythe and Wieschaus, 2016; Cho et al., 2022; Huang et al., 2021; Kemp et al., 2021). Although RD histone gene expression is activated during cycle 11, zygotic RD histone function is not needed until cycle 15, as the large stores of RD histone mRNA and protein deposited into the egg are sufficient for the embryo to complete all S phases through cycle 14 even in the absence of any histone genes (Gunesdogan et al., 2014; Smith et al., 1993). During G2 of cycle 14, maternal RD histone mRNA is degraded (Lanzotti et al., 2002), and subsequent cell cycles depend on new histone mRNA synthesis (Smith et al., 1993). We found that after the completion of S phase in cycle 14, RNA pol II remains localized to the HLB throughout G2 phase (Fig. 1B, Supplemental Video 2A-C). RNA pol II disperses at each mitosis when the HLB disassembles (White et al., 2007), followed by reaccumulating in HLBs as they reassemble after the completion of mitosis (Fig. 1C, Supplemental Video 1A-C, 2A-C)(Hur et al., 2020).

To further explore the regulation of RNA pol II dynamics at RD histone genes in early embryonic cycles, we visualized Spt6, a histone chaperone that binds directly to RNA pol II to promote rapid elongation through chromatin (Ehara et al., 2022; Filipovski et al., 2022; Miller et al., 2023; Vos et al., 2018). Live imaging using a protein trap line that expresses GFP-Spt6 revealed that Spt6 is concentrated in HLBs throughout S phase of cycles 12 and 13 (Fig. 1D, Supplemental Video 3A-C, 4A-C). Interestingly, we found that at ∼10 minutes into S_14_ Spt6 disperses from the HLB and is no longer enriched in HLBs during the remainder of S_14_ and most of G2_14_ (Fig. 1D). It then re-accumulates in the HLB very late in G2_14_, just before the cells enter mitosis (Fig. 1E, Supplemental Video 5). By contrast, RNA pol II remains concentrated in the HLB throughout all of S and G2 phase of cycle 14 (Fig. 1D, E, Supplemental Video 4A-C, 5A-C). Thus, RNA pol II and Spt6 are simultaneously enriched in HLBs during embryonic S phases when RD histone genes are expressed, but the regulation of their enrichment in HLBs differs.

### RNA pol II concentrates in HLBs independent of Cyclin E/Cdk2-mediated phosphorylation of Mxc

To explore the dynamics of HLB localization of RNA pol II in canonical 4 phase cell cycles we examined 3^rd^ instar larvae wing imaginal discs. This tissue is composed primarily of a single epithelium that arises from a group of ∼30 cells that begin proliferating after embryo hatching and expand in number during larval development to ∼35,000 cells via G1-S-G2-M cell cycles (Tripathi and Irvine, 2022) (Fig. 1A). We detected RNA pol II and HLBs by staining wing discs from 3^rd^ instar larvae homozygous for mCherry-Rpb1 with Mxc antibodies. We also stained these discs with the MPM-2 monoclonal antibody, which recognizes Mxc that has been phosphorylated by Cyclin E/Cdk2 (Calvi et al., 1998; White et al., 2011; White et al., 2007). In wing disc cells, RNA pol II is present throughout the nucleus and is also enriched in HLBs irrespective of whether they contained phospho-Mxc (P-Mxc) (Fig. 2A). We used Imaris software to segment the data using the anti-Mxc signal to identify HLBs, and then quantified the total amount of RNA pol II (mCherry-Rpb1 signal) and P-Mxc (MPM-2 signal) within thousands of HLBs from multiple wing discs by summing the pixel intensity values for individual channels (i.e. RNA pol II signal or MPM-2 signal) within each segmented HLB (Fig. 2B-D). For every wing disc, the set of values from each channel were normalized relative to the highest value of that channel within any HLB. This approach allows comparison of relative signal values within individual HLBs across the different channels and different wing discs.

**Figure 2.**
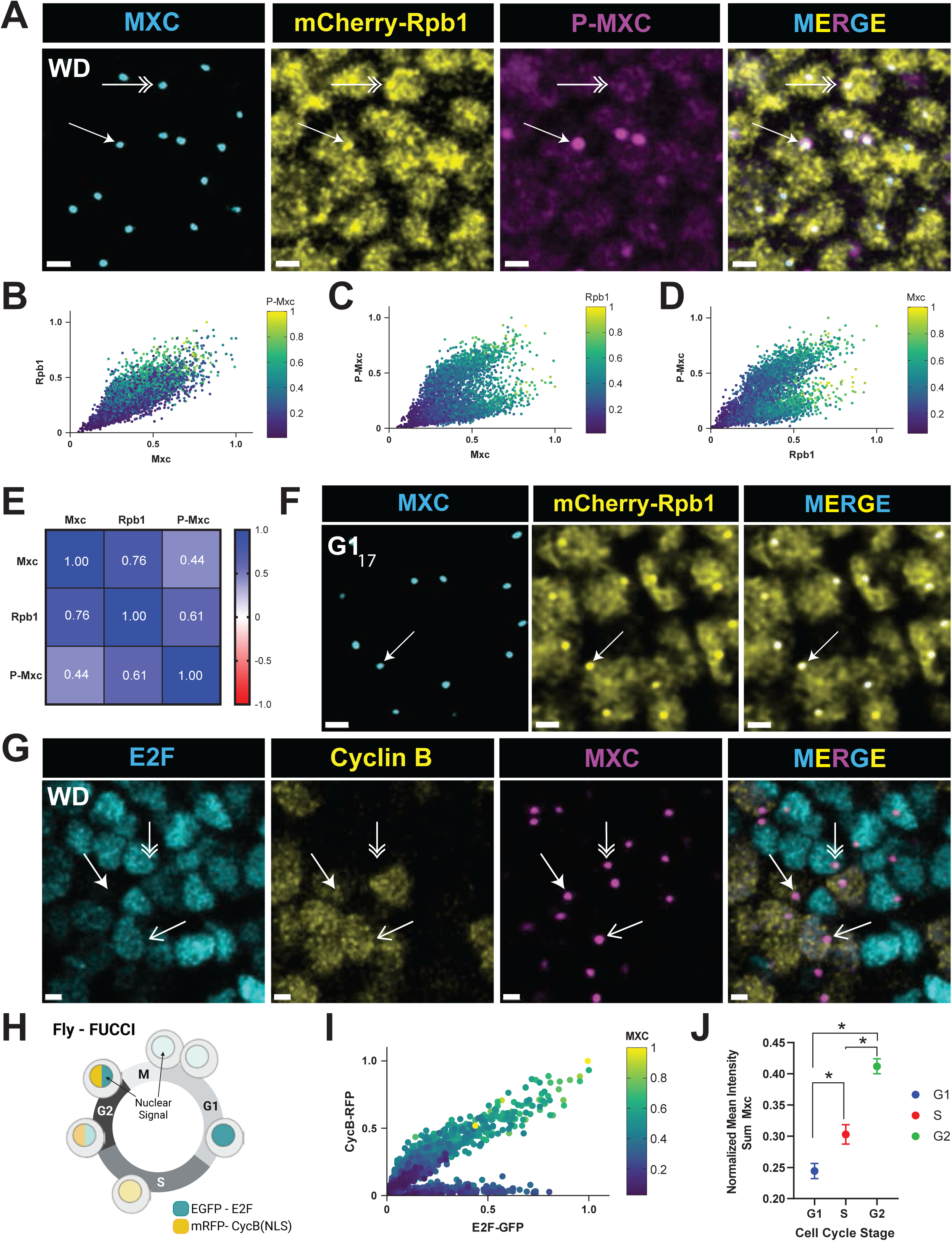
RNA polymerase II but not Spt6 is always enriched in HLBs **A)** Confocal micrographs of mCherry-Rpb1 wing disc cells from 3^rd^ instar larvae stained with anti-Mxc, anti-RFP, and MPM-2 (P-Mxc) antibodies to detect phospho-Mxc (P-MXC) in S phase. Single arrow indicates the HLB of a cell in S phase with active Cyclin E/Cdk2 (MPM-2 positive), and double arrow indicates the HLB of a cell not in S phase and without Cyclin E/Cdk2 activity (MPM-2 negative). Scale bar is 2 microns. **B-D)** Scatter plots of Mxc, RNA pol II, and MPM-2 signal intensities within 4390 HLBs from 1 wing disc as representatively shown in panel A. Similar results were obtained from 4 independent wing discs. Each dot represents an HLB segmented using the anti-Mxc signal, and each axis indicates normalized signal intensity measurements, with 1.0 being the highest. The values for MPM-2 (B), Rpb1 (C), and Mxc (D) are displayed as a heat map. **E)** Pearson correlation values of pairwise comparisons of signal intensity data from (B-D). **F)** G1_17_-arrested epidermal cells from germ band retracted mCherry-Rpb1 embryos stained with anti-Mxc and anti-RFP antibodies. All cells contain an HLB with enriched pol II (Arrow). Scale bars are 2 μM. **G)** Confocal micrographs of 3^rd^ instar larval wing discs ubiquitously expressing Fly-FUCCI (E2F-GFP, Cyclin B-RFP) and stained with anti-Mxc antibodies. Double arrow indicates an HLB in a G1 cell (E2F-GFP only), closed arrow indicates an HLB in an S-phase cell (low Cyclin B-RFP only), and the open arrow indicates an HLB in a G2 cell (E2F-GFP and Cyclin B-RFP). Note that not all nuclei show an obvious HLB because of the focal planes shown. Scale bar is 2 microns. **H)** Cartoon of cell cycle dependent expression of the Fly-FUCCI reporters. Made with BioRender. **I)** Scatter plot of E2F-GFP, Cyclin B-RFP, and Mxc signal intensities within 1405 HLBs from 1 wing disc, with each dot representing a single HLB. Similar results were obtained from 4 wing discs. Each axis indicates normalized signal intensity measurements, with 1.0 being the highest. The values for Mxc are displayed as a heat map. **J)** Plot of the mean normalized intensity sum of Mx in HLBs classified into G1, S, or G2 cell cycle phase using the Imaris machine learning function. Error bars are 95% confidence intervals for each class. Significance determined by Kruskal-Wallis test; **\*** = p< 0.0001.

Using this quantification method, we found that the amount of RNA pol II in HLBs was positively correlated (Pearson = 0.76) with the amount of Mxc (Fig. 2B, E). In contrast, the correlation between P-Mxc signal and either total Mxc (Fig. 2C, E) or total RNA pol II (Fig. 2D, E), revealed two distinct populations of HLBs, one with high phospho-Mxc (S phase) and one with low phospho-Mxc (G1 and G2 phase) (Fig. 2C, D). In addition, high levels of phospho-Mxc signal were apparent across a wide range of Mxc and RNA pol II amounts in HLBs (Fig. 2B, heat map). We conclude from these data that RNA pol II localizes to wing disc HLBs during S phase when RD histone genes are transcribed, as expected, but also in cells that are not MPM-2 positive and thus not in S phase. In contrast to RNA pol II, Spt6 only concentrates in HLBs that are P-Mxc positive (Supplemental Fig. 1A, open versed closed arrows). However, not all P-Mxc positive HLBs have an obvious accumulation of Spt6 (Supplemental Fig. 1A, double arrow), indicating that Spt6 is not enriched in HLBs throughout all of S phase. These data indicate that Spt6 is present in the HLB only during a portion of S phase in wing discs as we observed in embryonic S_14_.

To further explore the accumulation of RNA pol II outside of S phase, we examined RNA pol II in the epidermal cells during cycle 17 of embryogenesis. These cells are readily identified by embryonic stage and easily visualized because they are on the surface of the embryo. These cells do not have active Cyclin E/Cdk2 and are arrested in G1 phase (de Nooij et al., 1996; Lane et al., 1996). We found that in these G1-arrested cells, RNA pol II is enriched in HLBs even though they do not express RD histone genes (Fig. 2F and Supplemental Fig. 2). Thus, in both the embryo and wing disc, RNA pol II is enriched in the HLB outside of S phase and independently of Cyclin E/Cdk2 activity.

Another feature of the wing disc is the wide variation of Mxc and RNA pol II signal within HLBs. We considered whether this variation might reflect the increasing size of HLBs during interphase, an observation made previously in the syncytial nuclear cycles (Hur et al., 2020), as well as in human cells (Armstrong et al., 2023). To examine this issue in *Drosophila* wing discs, we employed a Fly-FUCCI line that expresses a fragment of the E2F1 protein and a fragment of the Cyclin B protein each fused to a different fluorescent protein (Zielke et al., 2014). These fusion proteins accumulate within the nucleus during different phases of the cell cycle, thereby providing a visual indication of G1 phase (Fig. 2G, double arrow), S phase (Fig. 2G, closed arrow) and G2 phase (Fig. 2G, open arrow) (Fig. 2H). We stained wing disc cells from the Fly-FUCCI line with Mxc antibodies and quantified the amount of Mxc signal versus the two reporters in segmented HLBs (Fig. 2I). We found that Mxc was lowest in G1 cells (Fig. 2I, dark blue circles) and highest in G2 cells (Fig. 2I, green and yellow circles). By training a machine learning algorithm from the Imaris software to classify thousands of cells as G1, S, and G2, we demonstrated that the mean intensity sum of Mxc in wing disc HLBs increases throughout interphase (Fig. 2J).

These results permit us to identify cell cycle phase by the level of Mxc in segmented HLBs. For instance, Spt6 was enriched in HLBs with intermediate levels of Mxc (Supplemental Fig. 1B), consistent with its recruitment only during S phase (Supplemental Fig. 1). In addition, HLBs with the highest levels of Mxc are in cells in G2 phase of the cell cycle, while the lowest levels would be in G1 cells.

### RNA pol II is necessary for HLB assembly

Our observations that RNA pol II is present in the HLB irrespective of S phase in amounts that correlate with the amount of Mxc were a surprise and raised the possibility that RNA pol II might contribute to HLB formation or growth. To determine if RNA pol II is necessary for HLB formation we used syncytial embryos since maternally supplied gene products mean that zygotic gene expression is not required prior to cycle 14 (Harrison et al., 2023). Thus, essential transcription machinery like RNA pol II can be removed without blocking development. However, gene products provided maternally also mean that we cannot inhibit gene function in syncytial embryos using zygotic mutant genotypes. Instead, we used a technique called “JabbaTrap” that sequesters GFP-tagged nuclear proteins on lipid droplets in the cytoplasm of syncytial embryos, thus removing their function from the nucleus (Seller et al., 2019). We performed the experiment in fully viable fly lines in which all molecules of either Mxc or a subunit of RNA Pol II was tagged with GFP (i.e. GFP-Mxc or GFP-Rpb3) while the other was tagged with a red fluorescent protein (i.e. Mxc-mScarlet or mCherry-Rpb1) (Cho et al., 2022; Kemp et al., 2021). By injecting RNA encoding JabbaTrap into live syncytial embryos from these lines we could deplete either Mxc or RNA pol II from the nucleus and monitor the ability of the other protein to form foci representative of HLBs.

When Mxc was depleted from syncytial nuclei using the JabbaTrap method, RNA pol II failed to form large foci (Fig. 3A). In these embryos smaller foci of RNA pol II were still apparent throughout the nucleus, indicative of transcription at other loci (Fig. 3A) (Guglielmi et al., 2013; Isogai et al., 2007). These data indicate that Mxc is required for the formation of large RNA pol II foci in syncytial nuclei, consistent with our previous work demonstrating that Mxc is necessary for both HLB formation and RD histone gene expression (White et al., 2011). Surprisingly, sequestering RNA pol II in the cytoplasm via JabbaTrap mRNA injection resulted in foci of Mxc-mScarlet that are smaller than in control embryos (Fig. 3B). This result indicates that Mxc can be recruited to RD histone genes when RNA pol II is depleted from the nucleus, but HLB growth is attenuated.

**Figure 3.**
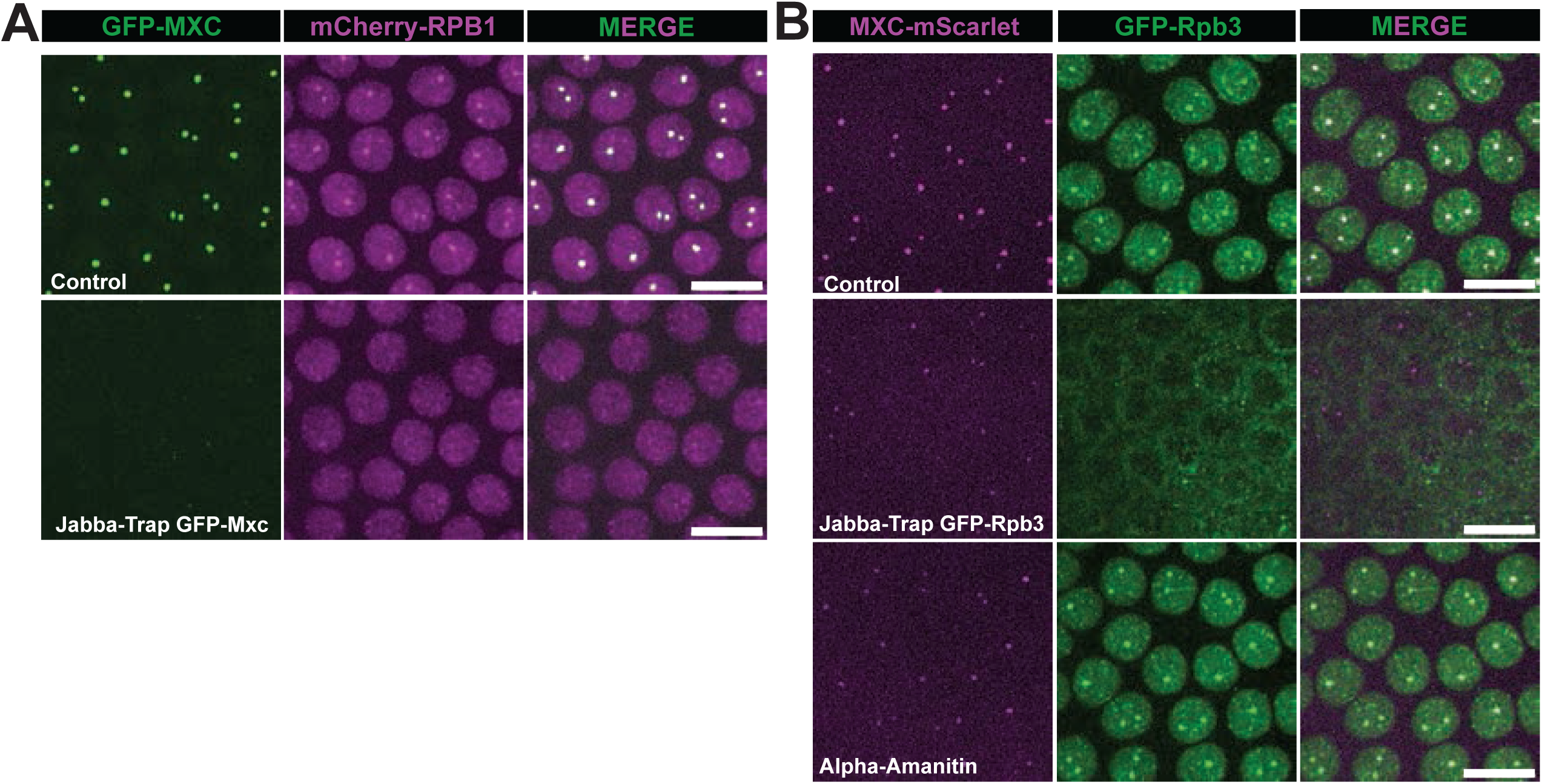
RNA pol II is necessary for HLB growth **A)** Single micrographs from live imaging of GFP-Mxc; mCherry-Rpb1 syncytial embryos injected with either water (top) or JabbaTrap mRNA (bottom). Note that cytoplasmic tethering of GFP-Mxc results in loss of large RNA pol II foci, likely from failure to form HLBs. **B)** Single micrographs from live imaging of Mxc-mScarlet; GFP-Rpb3 syncytial embryos injected with either water (top) or JabbaTrap mRNA (middle). Note that cytoplasmic tethering of RNA pol II results in smaller HLBs. (Bottom) Mxc-mScarlet; GFP-Rpb3 syncytial embryos injected with α-amanitin, which also results in smaller HLBs. Scale bars are 10 μM.

Injection of the RNA pol II inhibitor α-amanitin also resulted in smaller Mxc-mScarlet foci, consistent with previous observations (Fig. 3B) (Hur et al., 2020). Strikingly, however, α-amanitin did not prevent the formation of large RNA pol II foci, and these foci were coincident with the small Mxc-mScarlet foci (Fig. 3B). Since α-amanitin inhibits transcription by binding to RNA pol II that has engaged the DNA template and begun transcribing (Bushnell et al., 2002), this result is consistent with either RNA pol II recruitment and/or active transcription of the RD-histone genes making an important contribution to HLB growth (as judged by Mxc recruitment). Note that the α-amanitin results are also consistent with the possibility that RNAs expressed from loci other than histone genes may promote formation of the HLB.

### Transcription elongation at RD histone genes occurs when Cyclin E/Cdk2 is active

Next, we monitored nascent RD histone transcripts within the HLB to determine which steps in transcription are cell cycle regulated. To detect transcription of RD histone genes, we synthesized a pool of fluorescent oligonucleotide probes that hybridize to only the coding sequences (CDS) of all four core RD histone genes, which are transcriptionally co-regulated (Fig. 4A, Supplemental Fig. 3) (Salzler et al., 2013). We omitted the *H1* gene from the CDS probe because although its expression is replication-coupled, it utilizes different transcription factors (e.g. TRF2 versus TBP1) and is differently regulated than the core RD histone genes (Guglielmi et al., 2013; Isogai et al., 2007). There are ∼100 copies of each core RD histone gene at the *HisC* locus on each homologous chromosome. Furthermore, homologous chromosomes in *Drosophila* pair with one another during interphase, resulting in a single HLB in most cells (Fig. 4B, arrow). These features of genome organization in *Drosophila*, combined with the rapid export of processed mRNA(Adesnik and Darnell, 1972)^OBJ^ lets us visualize nascent RD histone gene transcription with high sensitivity.

**Figure 4.**
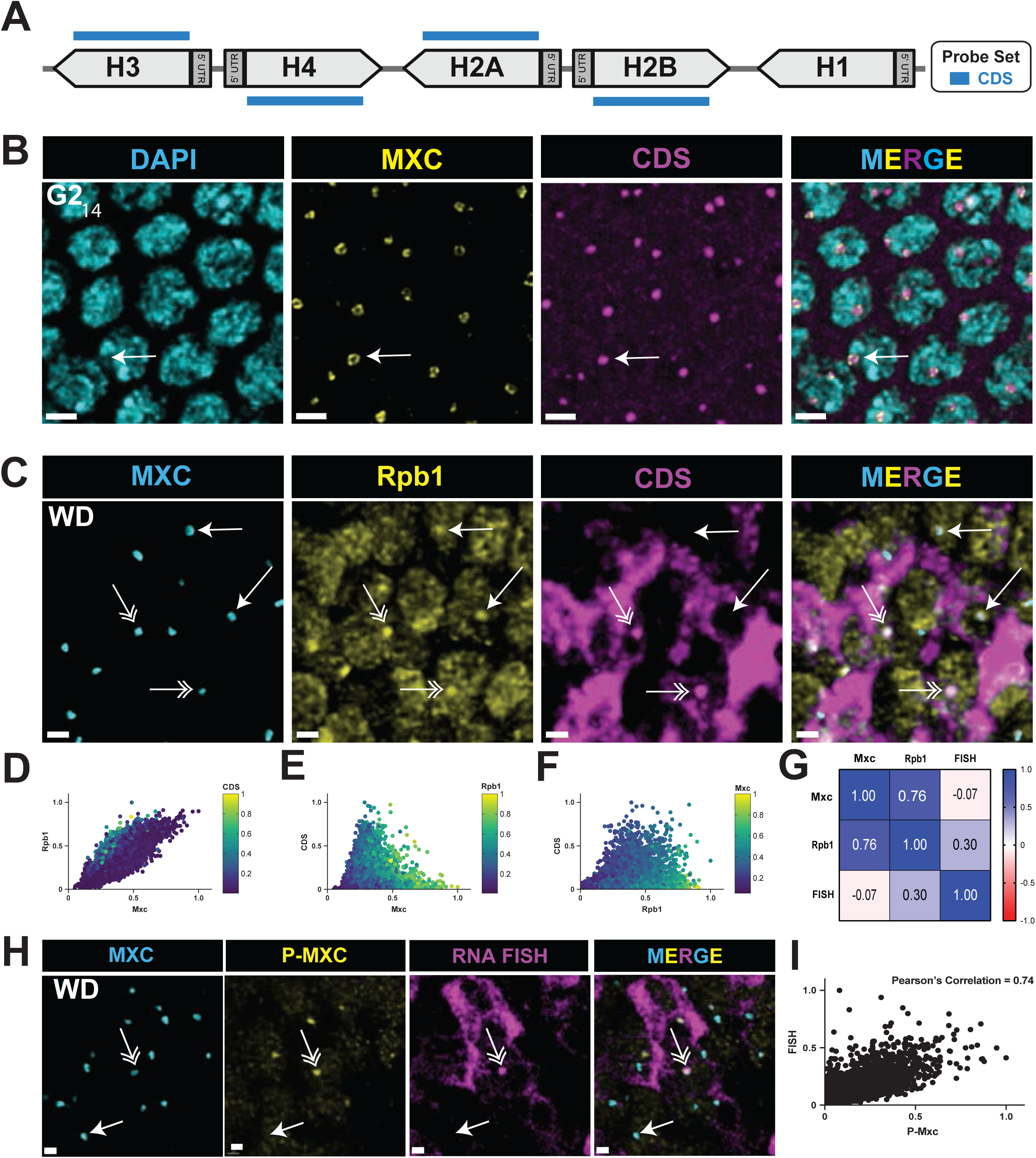
RNA polymerase II enrichment in HLBs occurs independent of S phase **A)** Diagram of the CDS Core RD-histone probe set which recognizes RNA transcribed from the coding region of the RD histone genes. Made with BioRender. **B)** Confocal micrographs of blastoderm embryos in G2 of cycle 14 hybridized with the CDS probe and stained with anti-Mxc antibodies and DAPI. The arrow indicates an HLB in a core shell configuration present in all HLBS at this stage with nascent RD histone RNA in the center. Nothe that there is little cytoplasmic RNA because the maternal mRNA is degraded at this stage. Scale bars are 3 μM. **C)** Confocal micrographs of mCherry-Rpb1 expressing wing disc cells hybridized with the CDS probe stained with anti-Mxc and anti-RFP (Rpb1) antibodies. RNA pol II and nascent RD histone transcripts are enriched in the HLB of S phase cells (double arrows), whereas only RNA pol II is enriched in the HLB of cells not in S phase (closed arrows). Note there is cytoplasmic histone mRNA in the S-phase cells. Scale bars are 1 μM. **D-F)** Scatter plots (as in Figure 2) of Mxc, RNA pol II (Rpb1) and nascent RD-histone RNA (FISH) signal intensities within 7061 HLBs from 1 wing disc representatively shown in (C). **G)** Pearson correlation values of pairwise comparisons of signal intensity data from (D-F). **H)** Confocal micrographs of wing disc cells hybridized with the CDS probe stained with anti-Mxc and MPM-2 (P-MXC) antibodies. Nascent RD histone transcripts are present in the HLB of S phase cells with active Cyclin E/Cdk2 (double arrows), but are absent in the HLB of cells not in S phase (closed arrows). Scale bars are 1 μM. **I)** Scatter plot of phospho-Mxc and nascent RD-histone RNA (FISH) signal intensities within 7749 HLBs from 1 wing disc representatively shown in (H). Similar results were obtained from 4 independent wing discs. Pearson correlation value of pairwise comparisons of signal intensity = 0.74.

We performed RNA FISH with embryos using the CDS probe and found that, like RNA pol II, nascent histone transcripts were present in HLBs throughout cycle 14, including during G2 phase (Fig. 4B). This result suggests that RNA pol II is actively synthesizing RD histone mRNA in G2_14_, and these cells contain active Cyclin E/Cdk2 and P-Mxc (Sauer et al., 1995; White et al., 2007). HLBs of G2_14_ embryonic nuclei appear in a “core-shell” configuration that we previously observed using super-resolution microscopy, with the nascent RD histone RNA in the center surrounded by Mxc (Kemp et al., 2021). In contrast, HLBs in cells not synthesizing RD histone mRNAs in the G1_17_-arrested epidermal cells lack this core-shell structure and have a more homogenous focus of both Mxc and RNA pol II (Supplemental Fig. 2A, B).

Using the CDS probe on wing imaginal discs we readily detected cells with nascent RD histone transcripts in HLBs containing RNA pol II (Fig. 4C, double arrows). We also observed cells without nascent transcripts in HLBs containing RNA pol II (Fig. 4C, single closed arrows). We quantified normalized signal intensities within wing disc HLBs (segmented by anti-Mxc signal) for Mxc, RNA pol II, and the CDS probe (Fig. 4D-F).

Each dot in Fig. 4D-F is the quantified signal from a single HLB assembled on paired homologous chromosomes and thus represents anywhere from 800 (i.e. G1) to 1600 (i.e. after *HisC* is replicated in S phase) core RD histone genes that likely behave asynchronously regarding transcription even when the HLB is “activated”. Consequently, the data provide a single cell view of the collective transcriptional behavior of these large gene clusters.

The quantification revealed that the relative amount of RNA pol II and Mxc were highly correlated and varied from relatively low levels of both proteins to relatively high levels of both proteins (Fig. 4D-G). In contrast, the nascent RD histone transcripts were not well correlated with the full range of either the Mxc or the RNA pol II signals. Rather, the nascent RD histone transcript signal using the CDS probe was highest within HLBs with intermediate amounts of Mxc and RNA pol II (Fig. 4E, F). As we showed in Fig. 2J, these HLBs are in cells in S phase. There was a strong correlation between P-Mxc levels and nascent RD histone transcripts within HLBs of wing discs hybridized with the CDS probe and stained with MPM-2 and anti-Mxc antibodies (Fig. 4H-I). These data indicate that high levels of nascent RD histone transcripts in cycling wing disc cells are present only in S-phase when Cyclin E/Cdk2 is active (i.e. MPM-2 positive), and that RNA pol II is also present in the HLB outside of S phase.

### In situ detection of transcriptional pausing at RD histone genes

Initiation of transcription by RNA pol II is immediately followed by co-transcriptional capping of the nascent transcript (Aoi and Shilatifard, 2023; Core and Adelman, 2019; Henriques et al., 2013) and subsequent recruitment of the negative elongation factor complex (NELF) and DRB sensitivity-inducing factor (DSIF), composed of Spt4 and Spt5. Transcription proceeds slowly for < 100 nts resulting in a “paused” RNA polymerase. Phosphorylation of Spt5, NELF and RNA pol II by P-TEFb kinase results in release of NELF and recruitment of Spt6 and PAF, resulting in activation of rapid elongation of transcription through chromatin (Aoi and Shilatifard, 2023; Chao and Price, 2001; Core and Adelman, 2019; Ni et al., 2008; Peterlin and Price, 2006).

Alternatively, the phosphatase activity of the Integrator complex can antagonize P-TEFb resulting in termination of the paused transcript, which is then rapidly degraded (Elrod et al., 2019; Huang et al., 2020; Stein et al., 2022; Wagner et al., 2023). The decision whether to enter rapid elongation or terminate a paused transcript contributes to the level of mature mRNA produced. Nascent transcripts associated with paused RNA pol II can be readily detected on *Drosophila* genes using genomic methods (Henriques et al., 2013; Nechaev et al., 2010), including on RD histone genes in cultured cells (Welch et al., 2015).

To test whether there were changes in pausing on the RD histone genes during the cell cycle, we synthesized two pools of differently colored fluorescent oligonucleotide probes that recognize either the 5’ or 3’ portions of the core RD histone transcripts (Fig. 5A, Supplemental Fig. 3). The 3’ probe set hybridizes to the last 200 nts of the mRNA located downstream of the RNA pol II pause site in these genes. Thus, this probe set detects nascent RD histone RNA associated with elongating RNA pol II and will detect all the transcripts detected with the CDS probe. We also designed a 5’ probe set to hybridize to the first ∼200 nts of RD histone RNAs including the ∼50 nt 5’ UTRs (Fig. 5A, Supplemental Fig. 3), which are absent from the CDS probe. The 5’ probe will detect nascent RD histone transcripts that are paused as well as those associated with elongating RNA pol II. The CDS probe will not readily detect paused transcripts since it lacks 5’UTR sequences. As with the CDS probe, we found that both the 5’ and 3’ probe sets detected nascent RD histone transcripts throughout embryonic cycle 14 (Fig. 5B, C). Thus, elongation occurs at RD histone genes during S_14_ and G2_14_ irrespective of whether Spt6 was enriched in the HLB (Fig. 1D, E). Using the 3’ probe we found that transcription elongation continues into the early stages of mitosis 14, with the nascent RD histone transcripts disappearing by metaphase as the HLB disassembles (Supplemental Fig. 4). Nascent elongating transcripts appear again within the HLB during S phase 15, which begins immediately after mitosis 14 (Supplemental Fig. 4).

**Figure 5.**
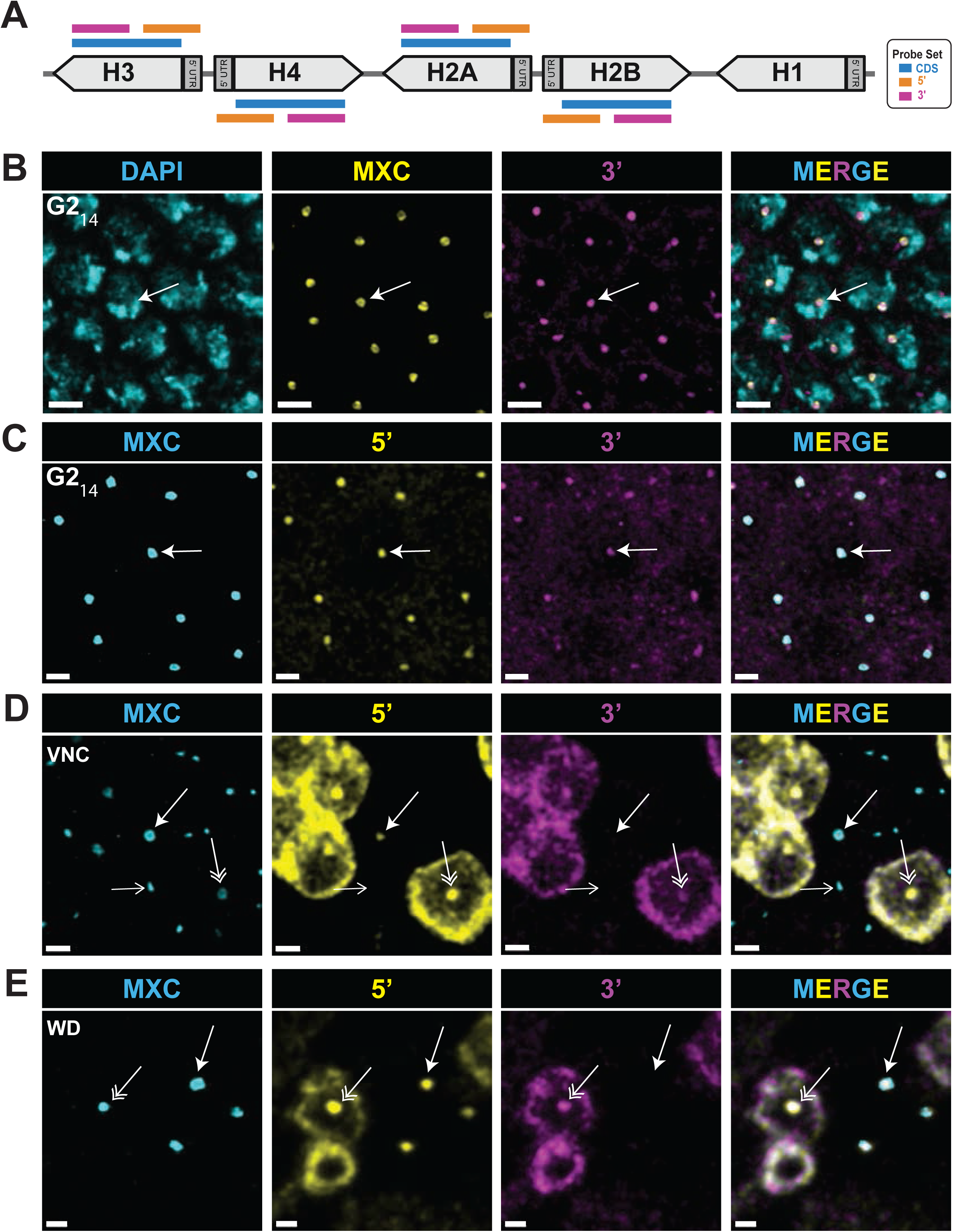
In situ detection of transcriptional pausing at RD histone genes **A)** Diagram of 5’ and 3’ Core RD-histone probe sets compared to the CDS RD-histone probe set (see Supplemental Fig. 3 for details). Made with BioRender. **B,C)** Confocal micrographs of embryonic cycle 14 cells in G2 phase hybridized with the 3’ probe and stained with anti-Mxc antibodies and DAPI (B) or both the 5’ and 3’ probes and Mxc (C). Nascent elongating transcripts are present in all cells (Arrow). Scale Bars are 3 μM in B and 2 μM in C. **D, E)** Confocal micrographs of cells in the ventral nerve cord of germ band retracted embryo (D) or wing imaginal disc cells (E) hybridized with both the 5’ and 3’ probes and stained with anti-Mxc antibodies. Double arrows indicate HLBs containing elongating nascent transcripts in an S phase cell, as noted by the presence of cytoplasmic RD histone mRNA. The closed arrows indicate HLBs with nascent RNA signal from the 5’ probe but not the 3’ probe, demonstrating paused transcripts. Note that these HLBs have a core shell configuration and that the cells are not in S phase as indicated by the lack of cytoplasmic RD histone mRNA. In D) the HLBs in the field that do not hybridize with either probe are epidermal cells arrested in G1, shown by the open arrow. Scale bars are 1 μM.

We found a different situation when examining cells within the ventral nerve cord of older embryos or cells from larval wing discs. In each of these tissues we detected HLBs simultaneously positive for both the 5’ probe and the 3’ probe within cells where we could unambiguously identify cytoplasmic RD histone mRNA as well (Fig. 5D, E, double arrows). In addition, we detected HLBs that were positive for the 5’ probe, but not the 3’ probe, and these cells did not contain cytoplasmic RD histone mRNA (Fig. 5D, E, closed arrows). These data indicate that in some cells RD histone gene transcription makes only paused transcripts which are then terminated. HLBs containing only the 5’ nascent transcript signal displayed a core shell configuration demonstrating that this structure is found associated with either paused or elongating transcripts at RD histone genes (Fig. 5D, E, closed arrow). HLBs arrested in G1_17_ lacked both the 5’ and 3’ nascent transcript signal and were not in the core-shell configuration (Fig. 5D, open arrow and Supplemental Fig. 2C), despite having high levels of RNA pol II (Fig. 2F). Thus, we detect three populations of HLBs regarding transcription of the RD histone genes: not initiated, paused, and elongating.

### Cell cycle-coupled pause-release controls RD histone gene expression

To address quantitatively whether transcriptional pausing at RD histone genes is cell cycle regulated, we stained larval wing imaginal discs with both the 5’ and 3’ probes and either Mxc (Fig. 6A-D) or EdU (Fig. 6E), or with the 5’ probe and both EdU and MPM-2 antibodies to mark P-Mxc in S phase cells with active Cyclin E/Cdk2 (Fig. 6F). We quantified the relative level of 5’ and 3’ nascent transcript signal within HLBs segmented using anti-Mxc antibodies (Fig. 6B-D). This analysis revealed two distinct populations of HLBs, one with high levels of both the 5’ and 3’ probes and one with high levels of only the 5’ probe (Fig. 6B). HLBs positive for both 5’ and 3’ probes were within cells that also contained cytoplasmic RD histone mRNA, and pulse labeling with EdU confirmed that these cells were in S phase (Fig. 6E, closed arrow). Cells with HLBs containing 5’ signal but lacking 3’ signal did not incorporate EdU and lacked P-Mxc signal (Fig. 6E, F, open arrows), which is found only in S phase cells that have active Cyclin E/Cdk2 (Fig. 6F, closed arrow). HLBs with intermediate levels of Mxc and thus from S phase (Fig. 2J) are the only ones expressing nascent elongating transcripts as detected by the 3’ probe (Fig. 6C, D). HLBs with the most Mxc and thus from G2 cells (Fig. 2J) had the highest amount of 5’ signal and low or no 3’ signal (Fig. 6B-D). These data indicate that in cycling wing imaginal disc cells RD histone transcription elongation occurs only during S phase, as expected to make mature histone mRNA for production of new histones, and that only paused transcripts are produced outside of S phase when Cyclin E/Cdk2 is not active.

**Figure 6.**
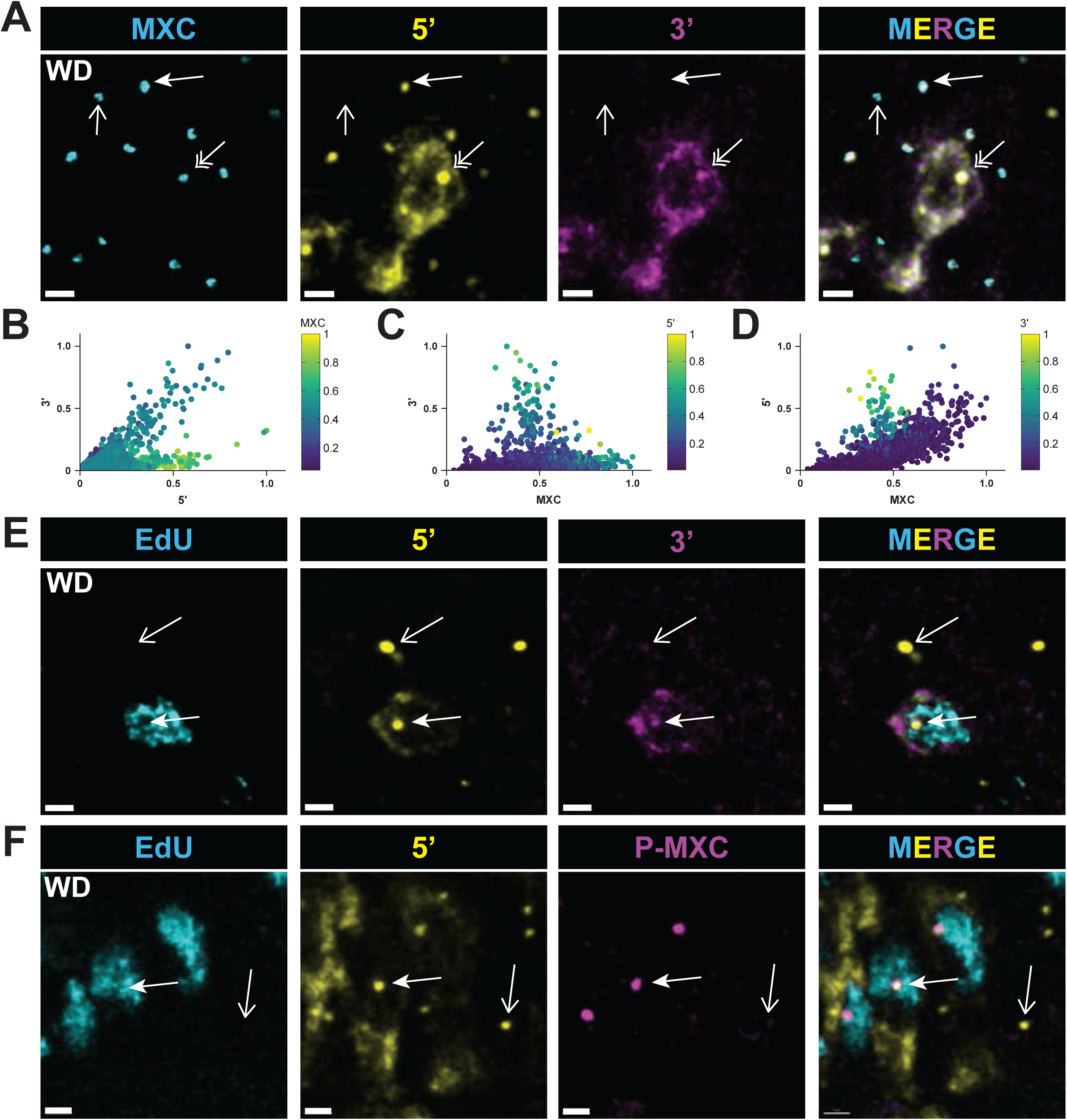
Release from transcriptional pausing at RD histone genes is cell cycle regulated **A)** Confocal micrographs of wing imaginal disc cells hybridized with both the 5’ and 3’ probes and stained with anti-Mxc antibodies. Double arrows indicate HLBs containing elongating nascent transcripts in an S phase cell. Closed arrows indicate HLBs with nascent RNA signal from the 5’ probe but not the 3’ probe, indicative of transcriptional pausing and no elongating transcripts in a cell not in S phase. Open arrows indicate an HLB lacking nascent RD histone transcripts, likely in a cell that has exited the cell cycle. Scale bars are 2 μM. **B-D)** Scatter plots of Mxc, 5’, and 3’ signal intensities within 1222 HLBs from 1 wing disc as representatively shown in panel A. Similar results were obtained from 4 independent wing discs. The values for Mxc (B), 5’ (C), and 3’ (D) signals are also displayed as a heat map. Note the clearly distinct population of HLBs with high 5’ signal but low 3’ signal. **E,F)** Confocal micrographs of wing imaginal disc cells pulse labeled with EdU and hybridized with the 5’ probe plus either the 3’ probe (E) or MPM-2 (P-MXC) antibodies (F). Closed arrows indicate HLBs in MPM-2 positive S phase cells, as indicated by extensive EdU labeling, and open arrows indicate an HLB in a cell not in S phase with only the 5’ probe and thus lacking elongating transcripts. Scale bars are 2 (E) and 1 (F) μM.

### Initiation of RD histone gene transcription is independent of Cyclin E/Cdk2

To determine directly whether RD histone gene transcription initiates in G1 prior to Cyclin E/Cdk2 phosphorylation of Mxc and then pauses, we analyzed eye imaginal discs. During the third larval stage this tissue undergoes a wave of differentiation associated with an apical constriction called the morphogenic furrow (MF) that progresses from posterior to anterior across the disc (Kumar, 2011). Cells within the MF are in G1. Immediately posterior to the MF, a subset of these cells begins differentiating into photoreceptor neurons while the remainder enter a synchronous S-phase called the “second mitotic wave”. This developmental program provides an opportunity to examine an easily identifiable G1-S transition in an endogenous tissue. We stained 3^rd^ instar larval eye imaginal discs with both the 5’ and 3’ probes and Mxc antibody to mark HLBs. HLBs of G1 cells within the MF (Fig. 7A ROI) contain 5’ nascent RD histone transcript signal but no 3’ signal (Fig. 7B), indicative of pausing of RD histone gene transcription during G1 prior to activation of Cyclin E/Cdk2 as cells enter S phase. Cells within S phase of the second mitotic wave have high levels of elongating nascent transcripts as well as cytoplasmic RD histone mRNA (Fig. 7A).

**Figure 7.**
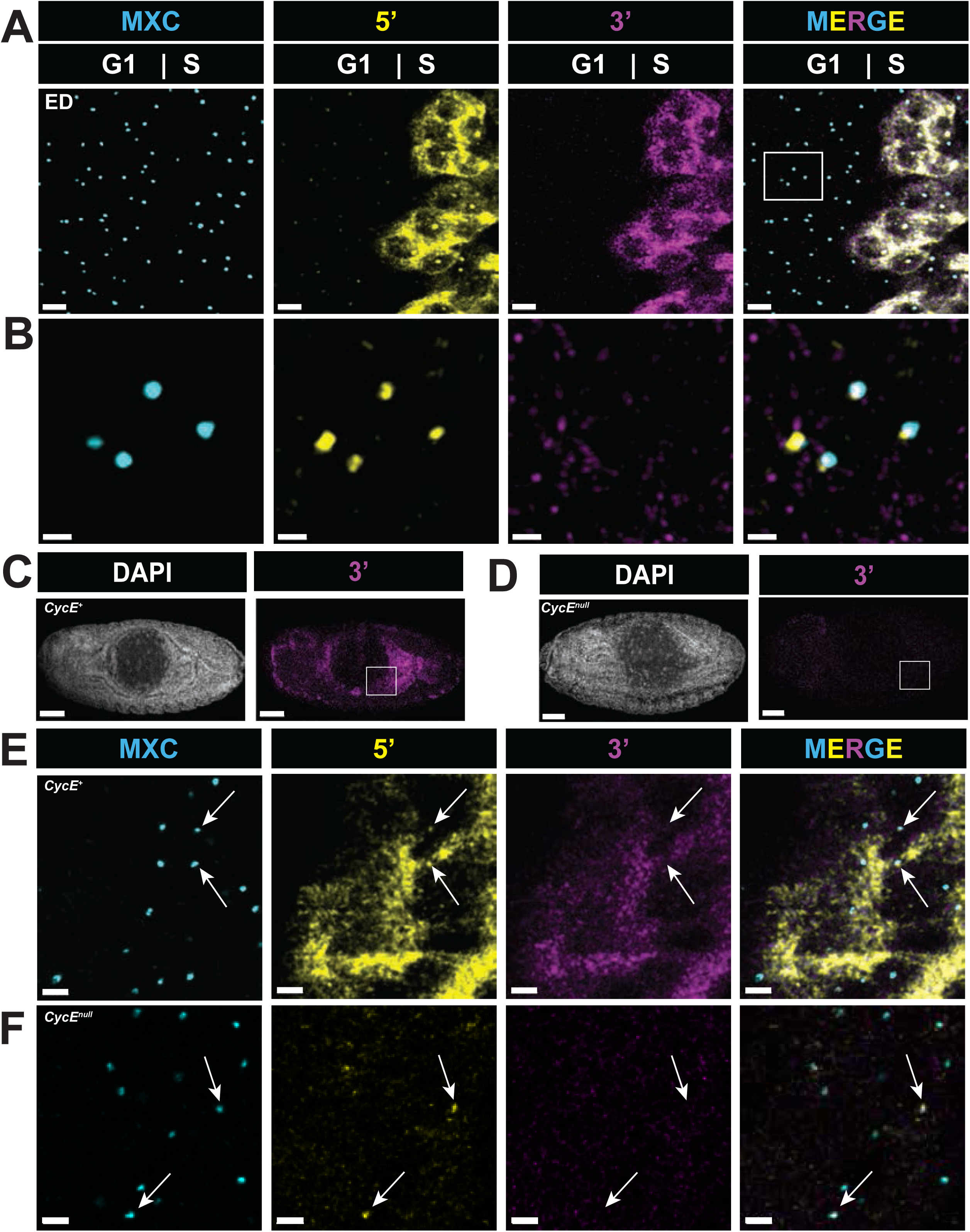
Transcriptional pausing at RD histone genes occurs independently of Cyclin E/Cdk2 **A)** Confocal micrographs of eye imaginal disc (ED) cells hybridized with both the 5’ and 3’ probes and stained with anti-Mxc antibodies. Scale bars are 3 μM. **B)** Boxed region from (A) of G1 cells that has been contrasted adjusted in both 5’ and 3’ channels showing HLBs with 5’ signal but lacking 3’ signal. Scale bars are 1 μM. **C,D)** Confocal micrographs of control sibling (*CycE^+^*) **(C)** and *Cyclin E* null mutant (*CycE^null^*) **(D)** Stage 14 embryos stained with DAPI and hybridized with the 3’ probe. Endocycling midgut cells in (D) contain cytoplasmic histone mRNA which is absent in (E) (boxed regions). Scale bar are 50 μM. **E,F)** Confocal micrographs of stage 14 embryonic anterior midgut cells (boxed regions in embryos from C and D) hybridized with both the 5’ and 3’ probes and stained with anti-Mxc antibodies in a control **(E)** and *Cyclin E* null mutant **(F)** embryo. Closed arrows indicate HLBs, which in *Cyclin E* mutants contain only 5’ probe. Scale bars are 2 μM.

To test whether initiation of RD histone gene transcription can occur independently of Cyclin E/Cdk2 activity, we analyzed *Cyclin E* null mutant embryos. In wild type embryos, tissues like the epidermis arrest in G1_17_, whereas others including the midgut enter endoreplication cycles to become polyploid and express high levels of RD histone mRNA (Fig. 7C, E). *Cyclin E* null mutant embryos develop normally until germ band retraction stages because of maternally supplied Cyclin E. However, in *Cyclin E* mutant embryonic midgut cells DNA replication does not occur and full-length RD histone mRNA fails to accumulate (Fig. 7D, F) (Lanzotti et al., 2004). We reasoned that developmental signals upstream of Cyclin E may still activate RD histone transcription in these cells but fail to activate the transition to elongating nascent 3’ RNA. Consistent with this hypothesis, we detected nascent 5’ nascent RD histone RNA signal in HLBs within *Cyclin E* mutant embryonic midgut cells but not 3’ signal or cytoplasmic RD histone mRNA (Fig 7F, arrows). These results indicate that RD histone gene transcription can initiate in the absence of Cyclin E/Cdk2 activity but cannot progress into active elongation and production of mRNA until Cyclin E/Cdk2 is activated to initiate DNA replication.

## Discussion

### RNA pol II recruitment to the RD histone locus contributes to HLB assembly but not cell cycle regulated RD histone gene expression

RD histone mRNA production is tightly coupled to S phase, and this coupling is critical for maintaining genome stability (Marzluff et al., 2008). However, despite many years of study, we still don’t understand all the mechanisms responsible for cell cycle-regulated RD histone mRNA production in vivo in animal tissues, particularly with respect to individual steps of the transcription cycle. The *Drosophila melanogaster* HLB provides several advantages in this regard, particularly the co-location of ∼800 co-regulated genes (i.e. ∼200 copies of *H3*, *H4*, *H2a*, and *H2b* in a diploid cell) and the associated proteins necessary for mRNA production within a large (∼1 μM in diameter) biomolecular condensate at a single locus (Duronio and Marzluff, 2017; Geisler et al., 2023). This nuclear organization allows relatively easy and highly sensitive microscopic detection of transcription. A prior analysis of cultured *Drosophila* S2 cells concluded that RNA pol II was present in the HLB only when RD histone genes were being transcribed during S phase (Guglielmi et al., 2013), suggesting that recruitment of RNA pol II to the HLB would provide a mechanism for controlling activation of RD histone gene expression. By contrast, we found using fluorescently labeled proteins expressed from endogenous genes that RNA pol II is concentrated within the HLB of G1_17_-arrested epidermal cells of the embryo without engaging in transcription and is present in the HLB of wing imaginal disc cells outside of S phase. Furthermore, preinitiation complex members TBP and TFIIA are present continuously in the HLB in cultured S2 cells (Guglielmi et al., 2013). Consequently, the regulated recruitment of RNA pol II to the HLB is not a mechanism for cell-cycle control of RD histone gene transcription in *Drosophila* tissues.

Our microscopic data cannot determine whether any or all the RNA pol II within the HLB is engaged with the DNA template, leaving open the possibility of different pools of RNA Pol II within the HLB (e.g. unbound and bound to DNA, Fig. 8). Given the large amount of RNA polymerase in the HLB, especially when cells are not synthesizing histone mRNA, there likely is always a pool of RNA pol II in the HLB that is not bound to DNA. A previous study using FRAP analysis concluded that in proliferating S2 cells in culture ∼80% of the RNA pol II molecules within the HLB were engaged in active elongation during S phase (Guglielmi et al., 2013). In *Drosophila* larval salivary glands up to 1/3 of the RNA pol II molecules at transcriptionally active *Hsp70* genes are not engaged in transcription (Versluis et al., 2024).

**Figure 8.**
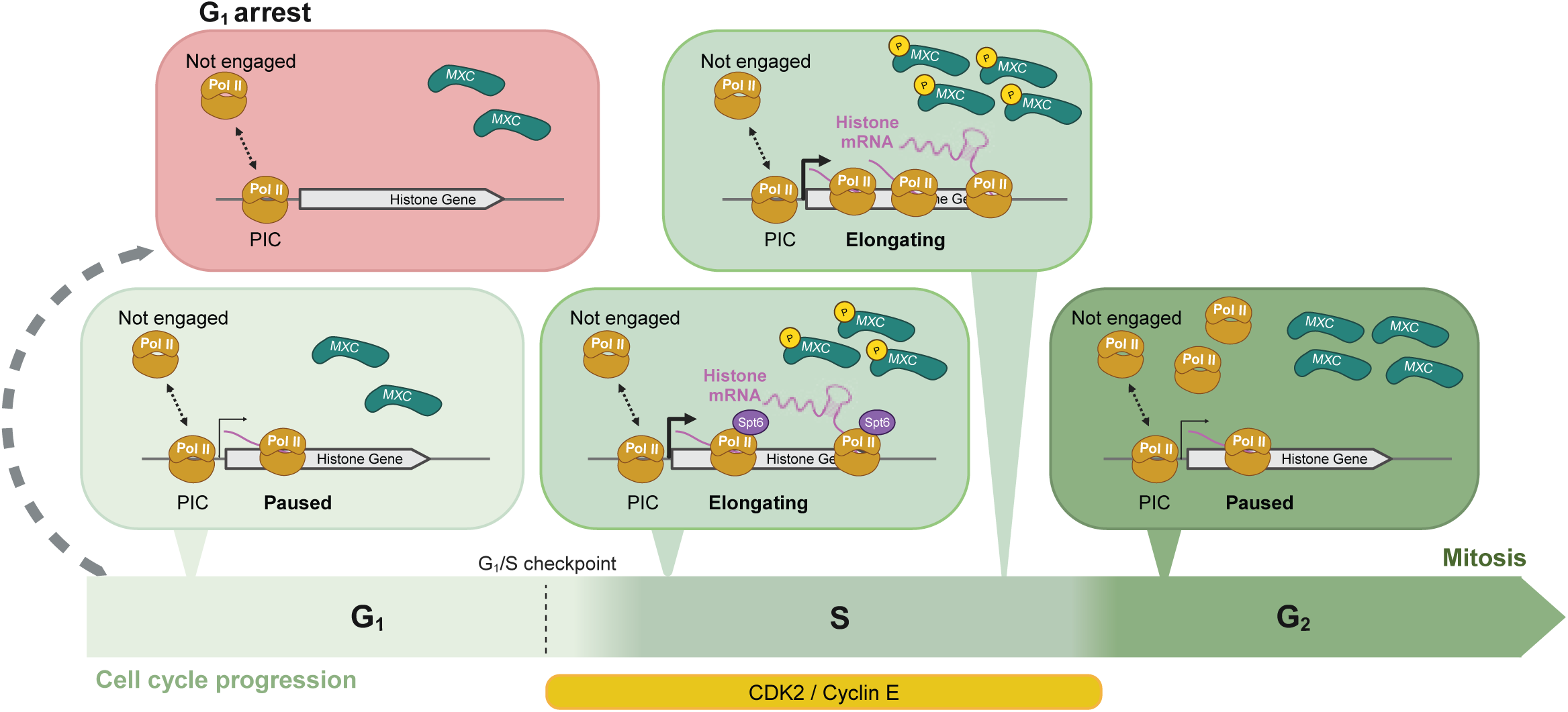
Summary of RD histone gene transcription Schematic representation of the presence of RNA pol II (Pol II), Mxc, P-Mxc, Spt6 in the HLB and RNA pol II transcriptional dynamics on the RD histone genes throughout the canonical cell cycle. Note G1 arrest is the only time where RNA pol II is not transcribing the RD histone genes, although it is still present in the HLB. Also note the amounts of both Pol II and Mxc in the HLB increase as the cell cycle progresses. Made with BioRender.

We found that depleting RNA pol II from nuclei in syncytial embryos resulted in smaller Mxc foci, indicating that HLB growth was attenuated and suggesting that RNA pol II is an integral component of HLBs. Here and previously we obtained similar results with α-amanitin injection in syncytial embryos (Hur et al., 2020). Furthermore, mutation of the *H3* and *H4* TATA boxes results in small HLBs at transgenic RD histone gene copies in larval salivary glands (Salzler et al., 2013). These data suggest that transcription at RD histone genes is necessary for HLBs to gain their typical size, but not for nucleation of Mxc biomolecular condensates, which happens stochastically in syncytial embryos lacking RD histone genes (Salzler et al., 2013). We don’t know whether HLB growth requires RNA pol II recruitment and/or binding to RD histone gene promoters, or actively transcribing RNA pol II and the presence of nascent histone mRNA, or conceivably a non-histone RNA transcribed at the locus or other loci. One possibility is that transcription increases accessibility of chromatin at the RD histone locus leading to a feed forward loop that recruits more Mxc and more RNA pol II.

Null mutations of *mxc* results in loss of HLB formation and loss of RD histone gene expression (Terzo et al., 2015; White et al., 2011). Accordingly, depletion of Mxc from syncytial embryos using JabbaTrap prevented the formation of large RNA pol II foci without affecting the formation of small RNA pol II foci throughout the rest of the nucleus. Mxc is a large (1800 aa) protein with a multimerization domain at its N-terminus essential for HLB assembly (Terzo et al., 2015) and a C-terminal domain that binds the pre-mRNA processing factor FLASH, as does human NPAT (Kemp et al., 2021; Yang et al., 2014). Mxc concentrates FLASH in the HLB, and this is essential for efficient cotranscriptional RD histone mRNA 3’ end formation (Tatomer et al., 2016).

Between its terminal domains Mxc is composed mostly of intrinsically disordered regions (IDRs) that could help recruit other factors to the HLB (DeRan et al., 2008; Miele et al., 2007; Zheng et al., 2003), such as RNA pol II, and which may contribute to the phase separation proprieties displayed by *Drosophila* HLBs (Hur et al., 2020).

Intriguingly, we observed large RNA pol II foci in α-amanitin treated syncytial embryos, even though these were associated with small Mxc foci. Components of the transcriptional machinery including RNA pol II form biomolecular condensates with properties consistent with phase separation (Boija et al., 2018; Cho et al., 2018; Cisse et al., 2013; Flores-Solis et al., 2023; Imada et al., 2021; Lu et al., 2018; Sabari et al., 2018). Thus, this observation suggests that a small Mxc condensate can nucleate a condensate of RNA pol II that can increase in size even though Mxc growth into a large condensate fails to occur.

### Cell cycle-coupled RD histone gene expression is mediated by transcriptional pausing

Cyclin E/Cdk2 is required for DNA replication as well as the accumulation of RD histone mRNA in *Drosophila* (Lanzotti et al., 2004). Cyclin E/Cdk2 phosphorylation of NPAT also activates RD histone gene expression during S-phase in mammalian cells (Armstrong et al., 2023; Armstrong and Spencer, 2021; Ma et al., 2000; Rogers et al., 2015; Wei et al., 2003; Ye et al., 2003; Zhao et al., 2000). Our results indicate that Cyclin E/Cdk2 stimulates RD histone gene expression by promoting the release of paused RNA pol II at RD histone genes resulting in production of histone mRNA. Previous single cell microscopic studies used probes lacking the 5’UTRs from RD histone mRNA and thus sequences contained in most of the paused transcripts.

Consequently, these studies did not detect transcription in G1 in cultured S2 cells (Guglielmi et al., 2013), or in G2 in embryos (Lanzotti et al., 2002; Lanzotti et al., 2004).

This function of Cyclin E/Cdk2 also explains the presence of elongating nascent transcripts in G2 phase of embryonic cycle 14 during embryogenesis. We found that RNA pol II is enriched in the HLB throughout G2_14_ and these cells continue to synthesize histone mRNA. During G2 in these cells Cyclin E/Cdk2 remains active and Mxc is phosphorylated (White et al., 2007). Cyclin E/Cdk2 remains active throughout the cell cycle up until late in interphase of cycle 16 when it is inactivated in the epidermis by developmentally controlled expression of the p21 Cdk inhibitor, Dacapo (de Nooij et al., 1996; Lane et al., 1996; Sauer et al., 1995). Consequently, epidermal cells enter G1 during interphase 17 and become quiescent. Here we observe no transcription of the histone genes, even though RNA pol II is present in the HLBs of these cells. Thus, transcription of paused RD histone gene transcripts is not dependent on Cyclin E/Cdk2, but Cyclin E/Cdk2 phosphorylation of Mxc (or potentially some other factors in the HLB) promotes release from transcriptional pausing and active elongation at RD histone genes. This is an important area for future investigation. Interestingly, there is one report that HIV-1 transcription is stimulated by CDK2 phosphorylation of P-TEFb (Breuer et al., 2012).

Promoter proximal pausing is a well described phenomenon in animal cells, including in *Drosophila* where the phenomenon and some of the regulatory factors were discovered (Marshall and Price, 1995; O’Brien and Lis, 1991). The presence of widespread proximal pausing was also first described in *Drosophila* (Nechaev et al., 2010). Release from transcriptional pausing provides a means for signal-mediated control of gene expression (Aoi and Shilatifard, 2023; Core and Adelman, 2019; Henriques et al., 2013). Notably, at *Drosophila Hsp70* genes paused transcripts are made and terminated constantly and production of mRNA only occurs after heat shock (Guertin et al., 2010). It is now recognized in both *Drosophila* and mammals that RNA pol II transcripts pause and either transition to elongation by P-TEFb or are terminated by the Integrator complex (Aoi and Shilatifard, 2023; Core and Adelman, 2019; Wagner et al., 2023). This decision point provides an additional important regulatory mechanism that controls the amount of mRNA synthesized from a particular gene (Boettiger and Levine, 2009; Lagha et al., 2013). Cell cycle-regulated pausing provides an opportunity for integration of different signals at different steps in mRNA synthesis. In addition, it provides an opportunity for assembly of factors onto the elongating RNA polymerase involved in subsequent pre-mRNA processing interactions. In a proliferating population of cells in growing tissues like *Drosophila* larval imaginal discs, release from transcriptional pausing on the histone genes by activation of Cyclin E/Cdk2 as cells enter S phase would be an effective way to achieve coordinated and rapid synthesis of large amounts of histone mRNAs encoded by multiple copies of the four core histone genes. Quiescent cells may not transcribe RD histone genes and prepare for reentry into the cell cycle in response to extracellular growth and/or developmental signals that in cooperation with specific transcription factors result in promoter proximal paused RNA pol II on the RD histone genes. These activated genes would not synthesize full length, mature histone mRNA, but would rapidly and coordinately do so when released from pausing upon Cyclin E/Cdk2 activation as cells transition to S phase. In this way S phase CDKs could provide a link to the cell cycle via a transcriptional “tuning” mechanism rather than a digital “on/off” switch, which would only be engaged when cells become quiescent (Core and Adelman, 2019). Further determining the molecular mechanisms of RD histone gene expression should contribute to our understanding of how cellular signaling networks couple transcriptional pausing and release to cell cycle progression.

## Materials and Methods

### Fly strains and husbandry

The following fly strains were used in this study. *D. melanogaster*: *y, w*, *mCherry-Rpb1, sfGFP-Mxc. w^1118^; If/CyO wg-lacZ; ub-GFP-E2F11-230^#5^ ub-mRFP1-NLS-CycB1-266^#12^/TM6B (BLM: 55124). y*, *w*, *mCherry-Rpb1, GFP-Spt6. y*, *w*, *Mxc-mScarlet, GFP-Spt6.* Oregon R. y,*w*, *Mxc-mScarlet*; EGFP-Rpb3*. w*; CycE^AR95^ cn^1^ pr^1^ bw^1^ wx^wxt^/CyO, P[ftz-lacB]E3 (BLM:6637). Cyclin E^AR95^/CyO wg-lacZ.* All recombinant chromosomes were first identified by detecting fluorescence larvae using a Leica fluorescent stereoscope and then confirmed via PCR. The following alleles have been described previously: mCherry-Rpb1 (Cho et al., 2022), EGFP-Rpb1 (Cho et al., 2022), Mxc-mScarlet (Kemp et al., 2021), sfGfp-Mxc (Hur et al., 2020), GFP-Spt6 (Buszczak et al., 2007), *Cyclin E^AR95^* (Duronio and O’Farrell, 1995).

### Live imaging and Immunofluorescence of fixed embryos

Embryos were collected and processed for imaging as described previously (Kemp et al., 2021). Live imaging was performed on a Leica SP8 imaging system using a 63x oil objective without any further digital zoom with the Lightning collection mode adjusted to equally balance speed and resolution for sfGFP-Mxc expressing embryos, or with just standard LSM collection for GFP-Spt6 expressing embryos. For time lapse images, time between collections were maximized for continual collection given the designated stack size for each individual embryo.

### Immunofluorescence of mCh-Rpb1, GFP-Spt6, and Fly-FUCCI larval tissues

Larval cuticles were inverted in PBS, fixed in 4% paraformaldehyde for 15 minutes, washed in PBS + 0.1% Triton x100 (PBST), and then stained with primary antibodies overnight at 4°C in 0.1% PBST. Primary antibodies were Millipore anti-MPM-2 Mouse IgG1 05-368 (1:10,000), Rockland anti-RFP rabbit 600-401-379 (1:1000), Rockland anti-GFP rabbit polyclonal 600-401-215 (1:1000), and anti-Mxc-C guinea pig (1:15,000) (White et al., 2011). Cuticles were then stained for 45 minutes at room temperature with secondary antibodies goat anti-rabbit Atto 488 (Sigma Cat #18772), goat anti-guinea pig Alexa 488 (Invitrogen), Goat anti-Rabbit IgG Alexa+ Fluor 555 (Invitrogen A32732), Goat anti-Mouse IgG1 Alexa Fluor 555 (Invitrogen A-21127), or Goat anti-Mouse IgG1 Alexa Fluor 647 (Invitrogen A21240), each diluted at 1:1000 in 0.1% PBST. Cuticles were then washed in 0.1% PBST, stained with 4’,6-diamidino-2-phenylindole (DAPI) (2µg/mL) for 10 minutes, and washed again. Wing discs were fully dissected off the larval cuticles and mounted on glass slides with ∼10ul of ProLong™ Glass Antifade Mountant (Invitrogen P36980) with a glass coverslip. Slides were left to dry overnight at room temperature in the dark and then imaged on a Leica SP8 confocal microscope LIGHTNING system.

### Immunofluorescence and RNA-FISH of mCh-Rpb1 larval tissues

To combine RNA-FISH with immunofluorescence, larval cuticles were inverted and stained as described above. After completion of secondary staining, cuticles were washed in 0.1% PBST, fixed in 4% paraformaldehyde for 5 minutes, and then washed in 0.3% PBST. Cuticles were then transitioned to Wash Buffer A (2x SSC, 10% Formamide (Sigma, 75-12-7), DEPC-Treated Water (Thermo, 4387937)) by washes in a dilution series of 4:1, 1:1 and 1:4 0.3% PBST: Wash Buffer A. Cuticles were then washed twice in Wash Buffer A for 5 minutes each at 37°C on a nutator. Prewarmed (37°C) Hybridization buffer (10% Dextran Sulfate (Fisher, 9011-18-1), 2x SSC, 10% Formamide (Sigma, 75-12-7), DEPC-Treated Water (Thermo, 4387937)) was added to cuticles and allowed to settle for 5 minutes. The CDS Core RD-histone Stellaris RNA-FISH probe set (LGC Biosearch Technologies) diluted to 100nM in Hybridization buffer was added and incubated overnight while nutating at 37°C. The following day hybridization buffer was removed, and cuticles were washed twice with prewarmed (37°C) Wash Buffer A for 30 minutes. Cuticles were then washed in 0.1% PBST, stained with 4’,6-diamidino-2-phenylindole (DAPI) (2µg/mL) for 10 minutes and washed again. Wing discs were fully dissected off the larval cuticles and mounted on glass slides with ∼10ul of ProLong™ Glass Antifade Mountant (Invitrogen P36980) with a glass coverslip. Slides were left to dry overnight at room temperature in the dark and then imaged on a Leica SP8 confocal microscope with LIGHTNING system.

### Immunofluorescence and RNA-FISH of embryos

Embryos were collected and stained as described previously (Kemp et al., 2021). After completion of secondary staining, cuticles were washed in 0.1% PBST, fixed in 4% paraformaldehyde for 5 minutes, and then washed in 0.1% PBST. Embryos were then washed in 50:50 0.1% PBST: Wash Buffer A for 10 minutes, followed by two five minute washes in Wash Buffer A at RT. Hybridization buffer was added to embryos and allowed to settle for 5 minutes. It was then replaced with fresh Hybridization buffer and incubated at 37°C on a thermomixer at 800rpm for 2 hours. The CDS or 5’ and 3’ Core RD-histone Stellaris RNA-FISH probe sets (LGC Biosearch Technologies) diluted to 50nM in Hybridization buffer was added and incubated overnight while nutating at 37°C. The following day hybridization buffer was removed, and embryos were washed once with prewarmed (37°C) Hybridization buffer for 30 minutes and then three times in prewarmed (37°C) Wash Buffer A for 15 minutes each. Embryos were then washed for 15 minutes in RT Wash Buffer A, then then washed in 0.1% PBST, stained with 4’,6-diamidino-2-phenylindole (DAPI) (2µg/mL) for 10 minutes and washed again. Embryos were mounted on glass slides with ∼10ul of ProLong™ Glass Antifade Mountant (Invitrogen P36980) with a glass coverslip. Slides were left to dry overnight at room temperature in the dark and then imaged on a Leica SP8 confocal microscope with LIGHTNING system.

### 5-ethynyl-2’-deoxyuridine (EdU) labeling, Immunofluorescence, and RNA-FISH of OreR larval wing and eye tissues

Larval cuticles were dissected and inverted in room temperature Grace’s Insect Medium (Gibco 11605094) and then nutated in Grace’s Insect Medium containing 0.25 mg/ml 5-ethynyl-2’-deoxyuridine (EdU) (Invitrogen A10044) for 30 min at room temperature to label S-phase cells. Cuticles were then washed once in Grace’s Insect Media and then fixed in 4% paraformaldehyde for 15 minutes, washed in PBS + 0.3% Triton x100 (0.3% PBST) 3 times for 10 minutes. They were then washed for 30 minutes in 3% BSA (Sigma # a4503). To detect EdU, the cuticles were incubated with Alexa Fluor™ 488 Azide (Alexa Fluor™ 488 5-Carboxamido-(6-Azidohexanyl) (Invitrogen A10266) and click detection reagents (Click-iT™ Cell Reaction Buffer Kit; Invitrogen C10269) at room temperature for 30 min to label EdU via a copper (I) catalyzed click reaction. Cuticles were then washed twice for 15 minutes in 3% BSA, in 0.1% PBST, quick fixed in in 4% paraformaldehyde for 5 minutes, and washed again in 0.1% PBST. To combine with immunofluorescence and RNA-FISH larval cuticles were then stained as described above using the 5’ Core RD-histone and 3’ Core RD-histone Stellaris RNA-FISH probe sets (LGC Biosearch Technologies). All Oregon R larval wing and eye tissues stained with one or both 5’ Core RD-histone and 3’ Core RD-histone probe sets underwent this full protocol, including when replicating DNA was not being detected by leaving EdU out of the culture medium (this improved detection of FISH signal likely because of the additional wash steps associated with the EdU labeling protocol).

### Generation of RNA Fluorescent in situ hybridization probes

Custom Stellaris RNA-FISH probes (Orjalo et al., 2011) targeting either the coding sequences (CDS Core RD-histone), the 5’ part of the mRNA sincluding the 5’ UTR (5’ Core RD-histone), or the 3’ part of the coding sequences (3’ Core RD-histone) of the core *D. melanogaster* histone genes (*H2A*, *H2B*, *H3*, *H4*) were generated using the Stellaris® RNA-FISH Probe Designer (LGC Biosearch Technologies, Petaluma, CA) and labeled with either Quasar 570 or Quasar 670 (Supplemental Fig. 3). The oligonucleotides used for each probe set are listed in the following table.

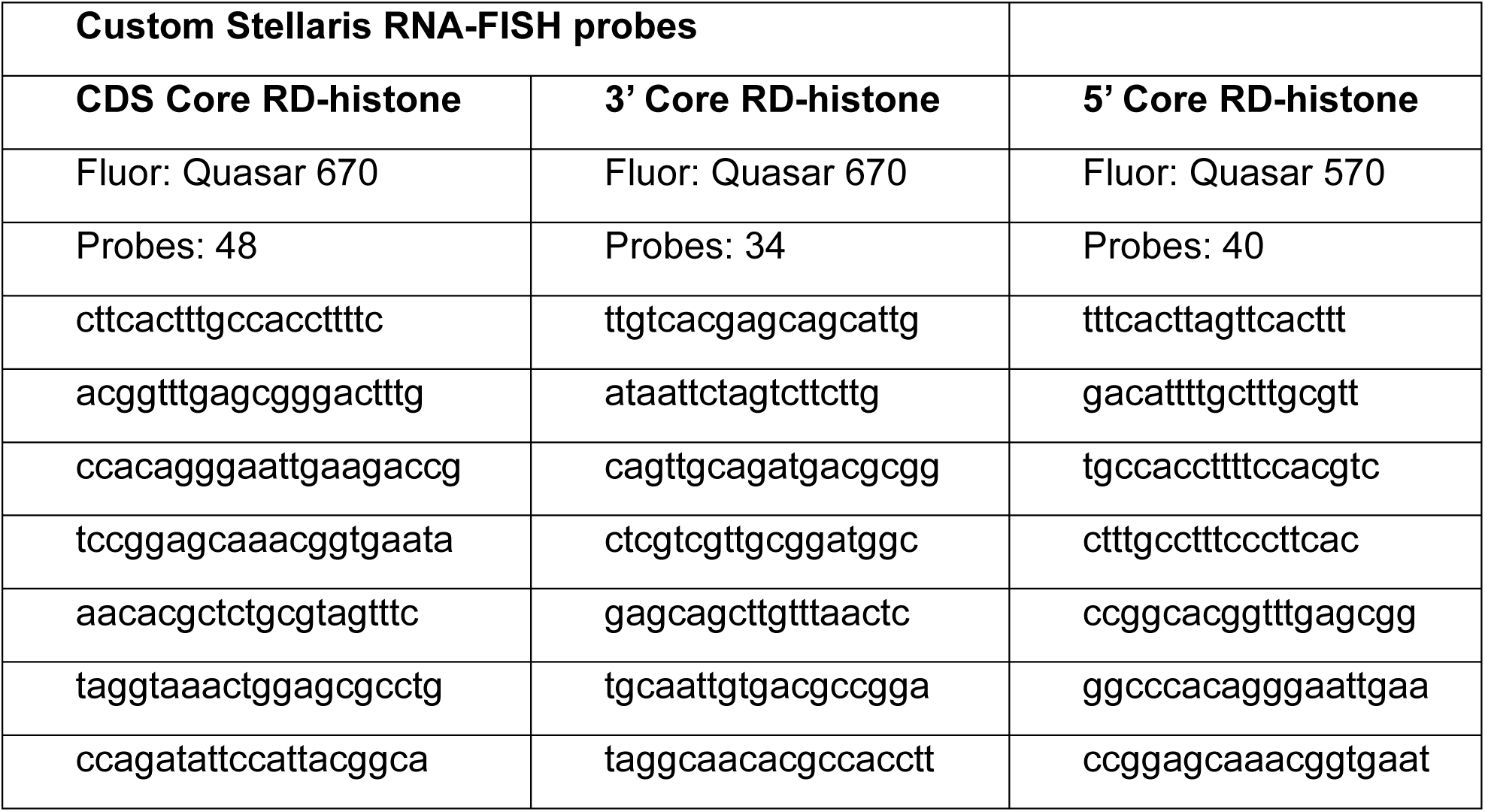
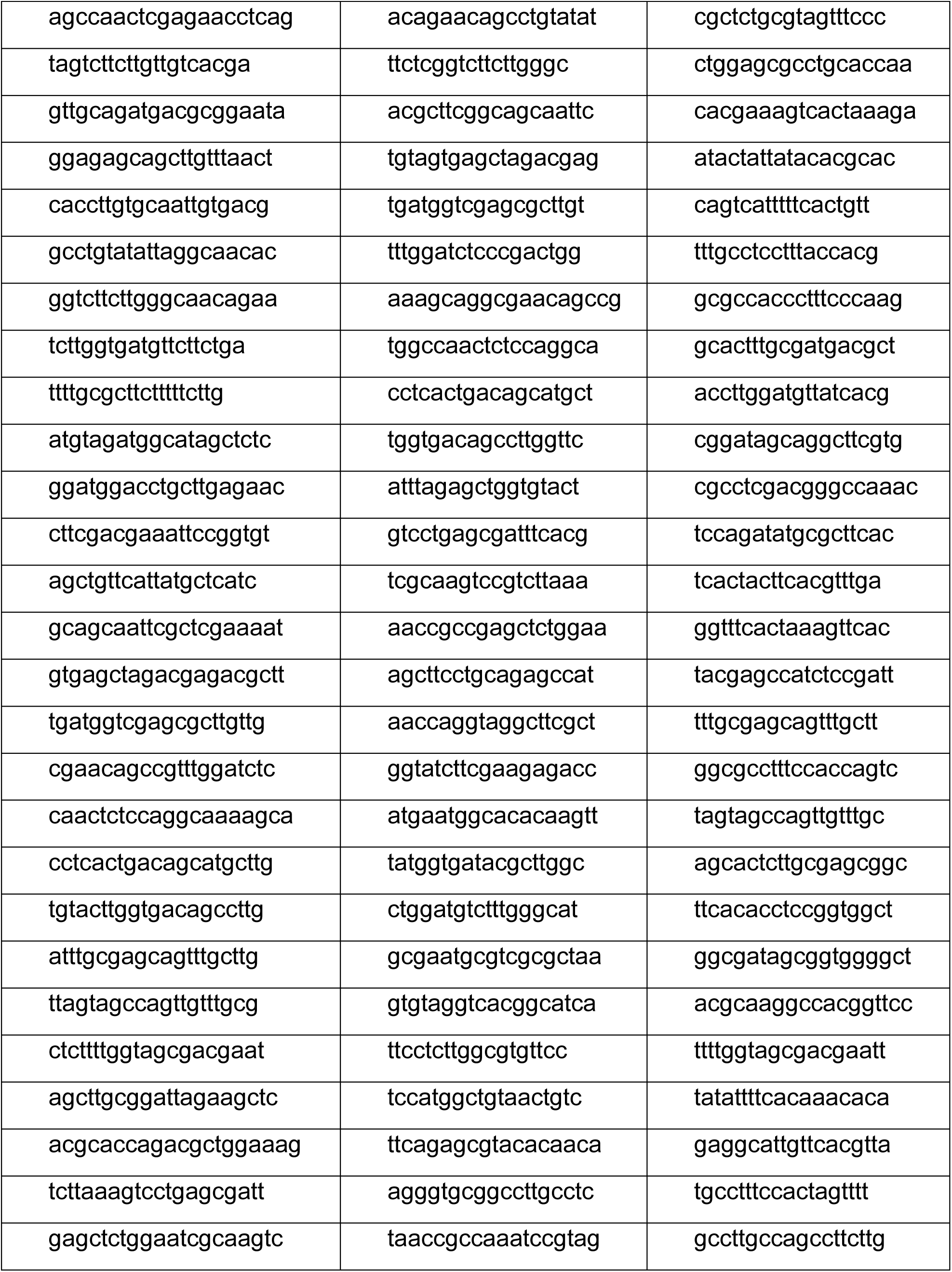
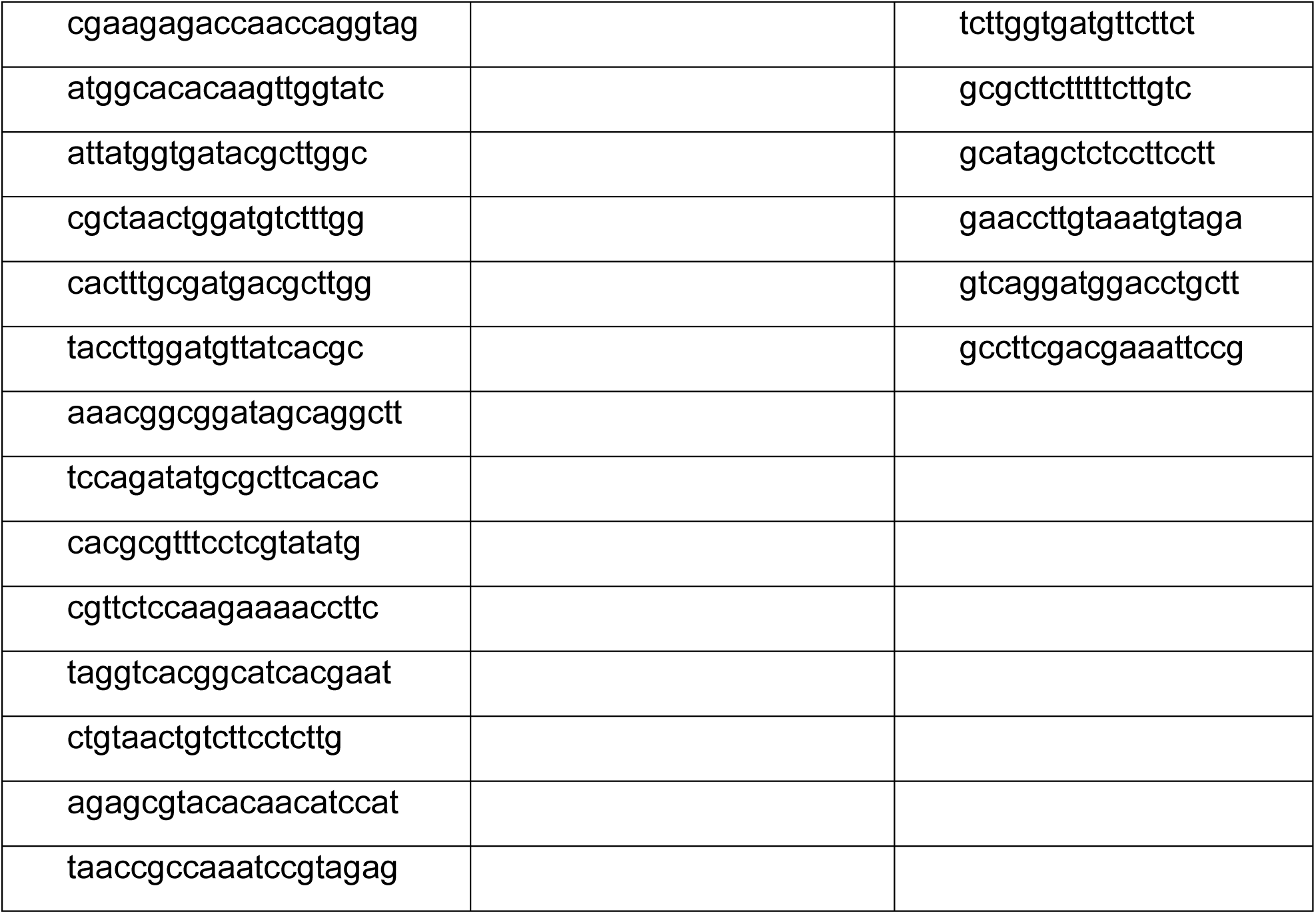

### Image Analysis and Quantification

All images were analyzed using FIJI and IMARIS software. To determine the levels of different factors present in the HLB, individual HLBs were segmented using Mxc as the source channel in the IMARIS spot function. For Mxc spots, background subtracted Mxc was used as the source channel for segmenting spots with an estimated XY diameter = 0.534 µm and estimated z diameter = 1.07 µm but with allowance for growing regions. Modeling of PSF-elongation along the z-axis was also used. Spots were filtered based on the intensity sum of square of Mxc with the threshold being set automatically but adjusted by eye for each image to maximize the HLBs capture while excluding artifacts. Object statistics were then exported and analyzed.

For classification of cell cycle stage, HLBs were segmented using Mxc as above and then classified using the spots machine learning module. The model was trained on n=30 HLBs on one image and then retrained twice more by correcting HLBs mis-classified into the wrong cell cycle stage. Once all spots in the training image were correctly classified, the model was applied to four new images, and the classes and statistics were exported and analyzed as described below.

### Analysis of quantification

The scale of Intensity sum values varied not only from channel to channel but from wing disc to wing disc within the same sample and under the same imaging conditions. To allow comparison between different discs and channels, values were normalized by dividing each spot’s intensity sum value from each channel by the max value within a dot of that channel, resulting in a value from 0 to 1 (i.e. Disc 1, Spot #1 Normalized value = Disc 1, Spot #1 ch.2 intensity sum/ Disc 1 Max ch.2 intensity sum). These normalized values were then plotted using GraphPad Prism.

### Fixed JabbaTrap Experiments

For the GFP-Spt6 JabbaTrap experiments we constructed lines with GFP-Spt6 and either Matα-Tub-Gal4 on the second chromosome or with UASp-JabbaTrap-Bicoid3’UTR on the third chromosome. These two stocks were then crossed to create females that were homozygous for GFP-Spt6 and heterozygous for Matα-Tub-Gal4 and UASp-JabbaTrap-Bicoid3’UTR. These females were then placed into cages with apple juice plates with GFP-Spt6; Uasp-JabbaTrap-Bicoid3’UTR males. After 24 hrs embryos 1-3 hrs old were collected, fixed, and processed as described above.

### Live JabbaTrap Experiments

Young embryos (1-1.5 hour after egg laying) were collected, rinsed with water, dechorionated with 40% bleach, and then washed thoroughly with water. Embryos were then transferred to grape juice agar plates, aligned and transferred to coverslips with the glue derived from double-sided tapes using heptane, and then slightly dehydrated by incubating in a desication chamber. After covering with halocarbon oil (1:1 mixture of halocarbon oil 27 and 700), embryos were injected with 750 ng/μl JabbaTrap mRNA synthesized as described (Seller et al., 2019). Once JabbaTrap proteins accumulated sufficiently to sequester GFP-tagged target proteins (typically 10-20 minutes later), embryos entering cycle 13 were subjected to live imaging on an Olympus IX70 microscope equipped with PerkinElmer Ultraview Vox confocal system as described (Cho and O’Farrell, 2023).

## Acknowledgements

We are grateful to iBGS for supporting the Leica confocal microscope used throughout this study, and the Department of Biology Microscopy Core. This work was supported by NIH F31HD113267 to M.S.G., NIH R35GM136324 to P.H.O, NIH R35145258 to R.J.D and NIH R01GM58921 to W.F.M. and R.J.D.

---

**Supplemental Figure 1.**
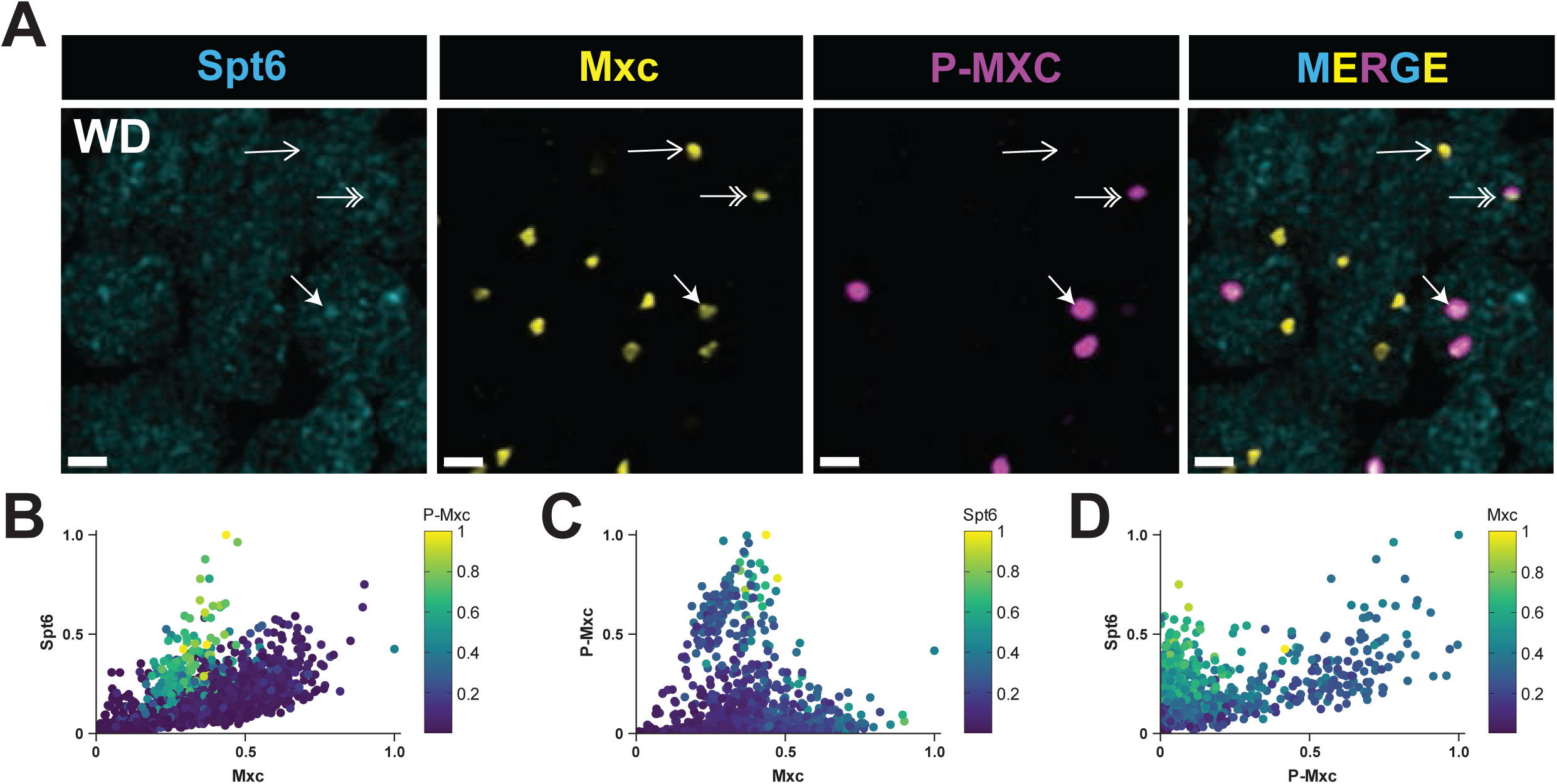
Spt6 concentrates in the HLB only during a portion of S-phase **A)** Wing disc cells from 3^rd^ instar larvae stained with anti-Mxc, anti-GFP (Spt6), and MPM-2 (P-Mxc) antibodies. Closed arrow indicates an HLB with enriched Spt6 in an S phase cell, whereas double arrow indicates an HLB lacking enriched Spt6 in an S phase cell. Open arrow indicates an HLB lacking Spt6 of a cell that is not in S phase. Scale bars are 1 μM **B-D)** Scatter plots of Mxc, Spt6, and phospho-Mxc (P-Mxc) levels within 1671 segmented HLBs from 1 wing disc as representatively shown in panel (A). Similar results were obtained from 4 independent wing discs. The values for P-Mxc (B), Spt6 (C), and Mxc (D) signals are also displayed as a heat map. Quantification of Spt6 and P-Mxc signal in segmented HLBs revealed three distinct populations of HLBs: 1) those that contain only Mxc, which are the most abundant (B, dark blue circles), 2) those that are high for both phospho-Mxc and Spt6 (C, yellow and green circles), and 3) those with high phospho-Mxc signal but low Spt6 signal (C, D).

**Supplemental Figure 2.**
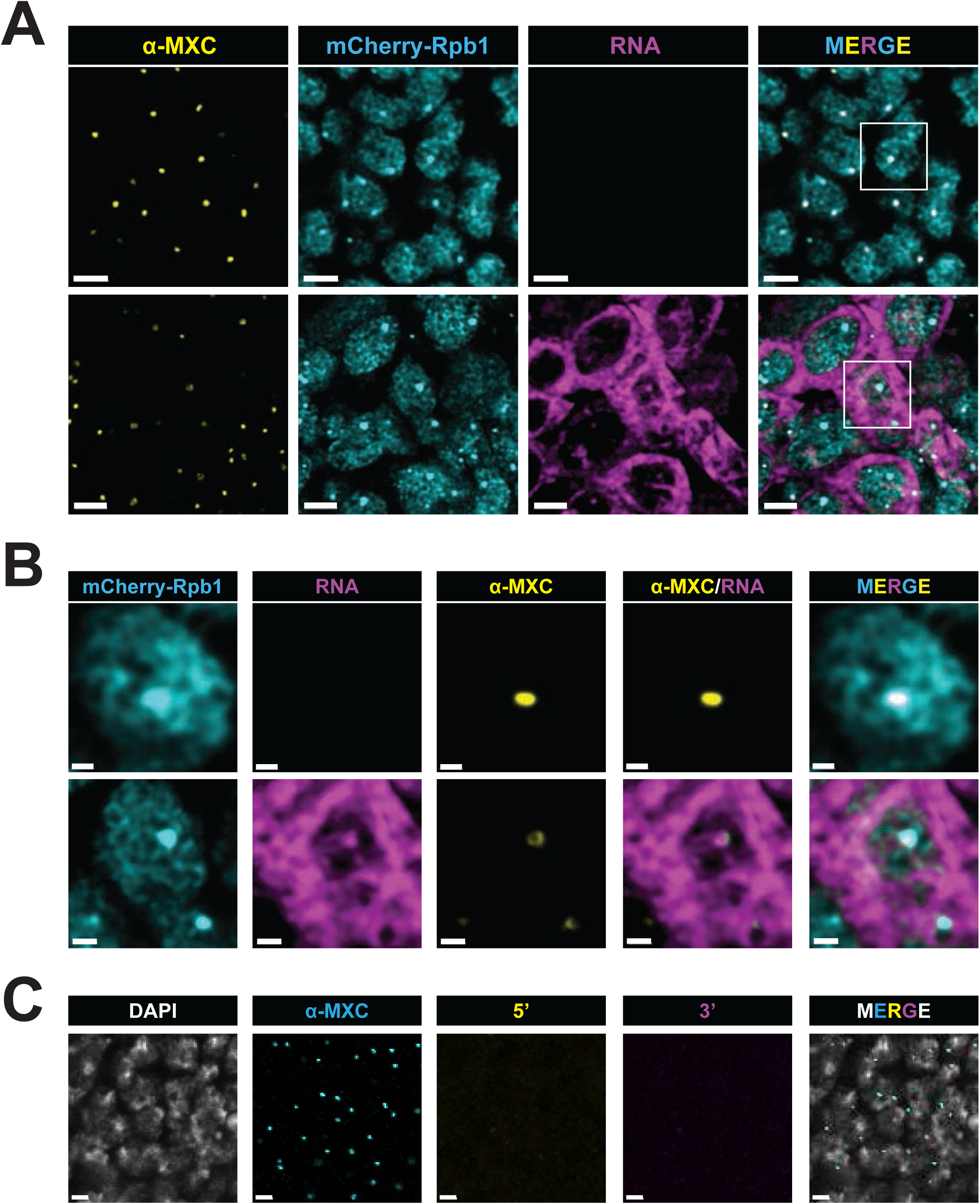
: **A)** Micrographs from a single representative mCherry-Rpb1 post stage 12 embryo hybridized with the CDS probe set and stained with anti-Mxc and anti-RFP antibodies. Epidermal cells arrested in G1_17_ are shown in the top panels, and a section of the VNC focused on replicating neuroblasts is shown in the bottom panels. Scale bars are 3 μM. **B)** Single nuclei from regions of interest from A. Note that we are unable to detect nascent RNA in epidermal cells (top panels) but detect both nascent and cytoplasmic RNA in the neuroblast (bottom panels). Also note the appearance of Mxc staining in the neuroblast and that the hole in the “donut” shape is filled with nascent RNA signal. Scale bars are 1 μM. **C)** Micrographs from a wildtype post stage 12 embryo hybridized with both the 5’ and 3’ probe set and stained with DAPI and anti-Mxc antibody showing epidermal cells arrested in G1_17_. These cells do not contain paused or elongating transcripts. Scale bars are 2 μM.

**Supplemental Figure 3.**
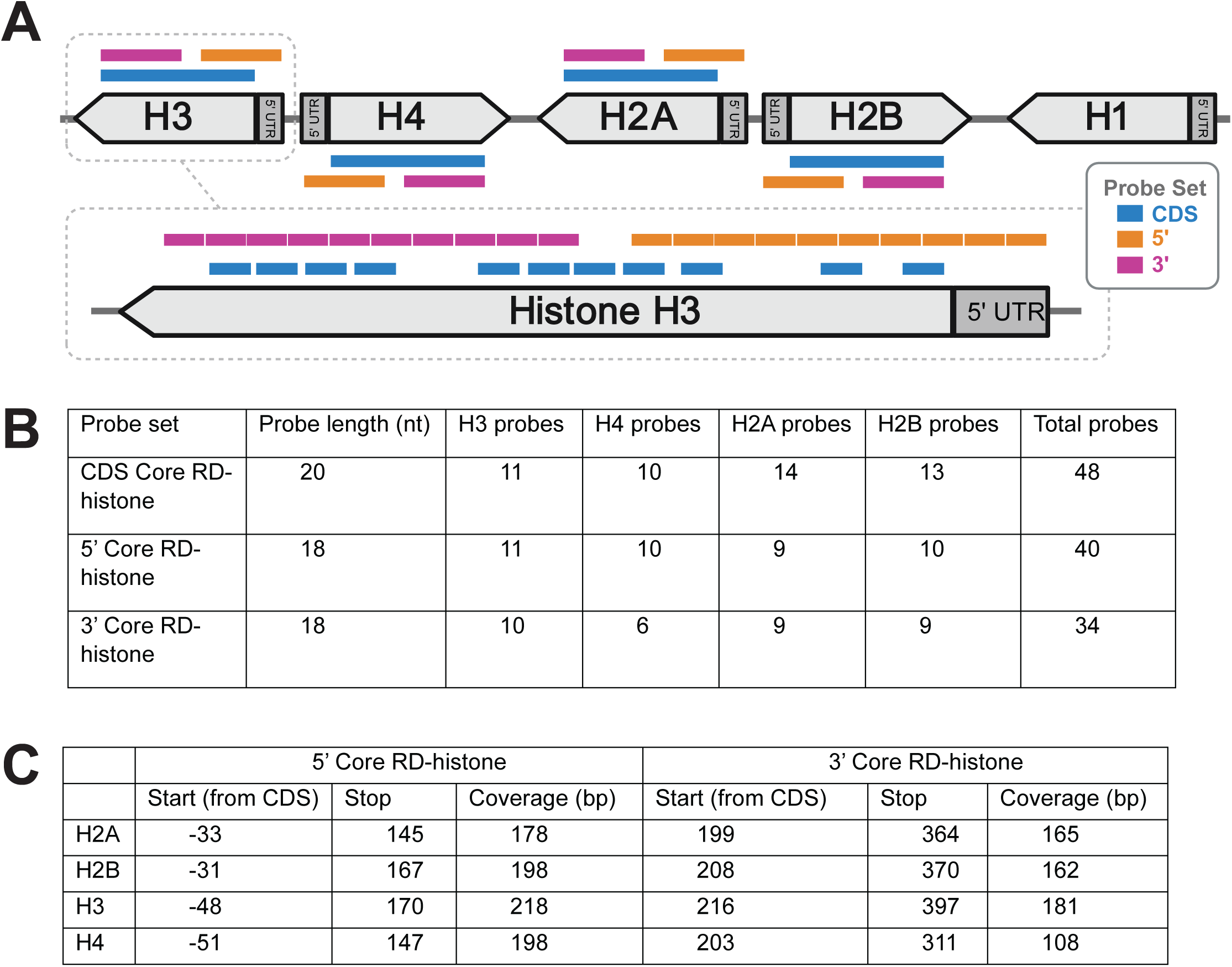
: **A)** Cartoon showing the coverage of the three RNA fish probe sets (CDS, 5’ and 3’) in all core replication dependent histone genes. Inset shows a zoomed in example of coverage for the *H3* gene showing the relative binding locations of individual probes from the different probe sets. Made with BioRender. **B)** Table outlining the probe lengths and number of probes per gene for each probe set. **C)** Table demonstrating the coverage and sequences used for each histone gene in the 5’ and 3’ FISH sets.

**Supplemental Figure 4.**
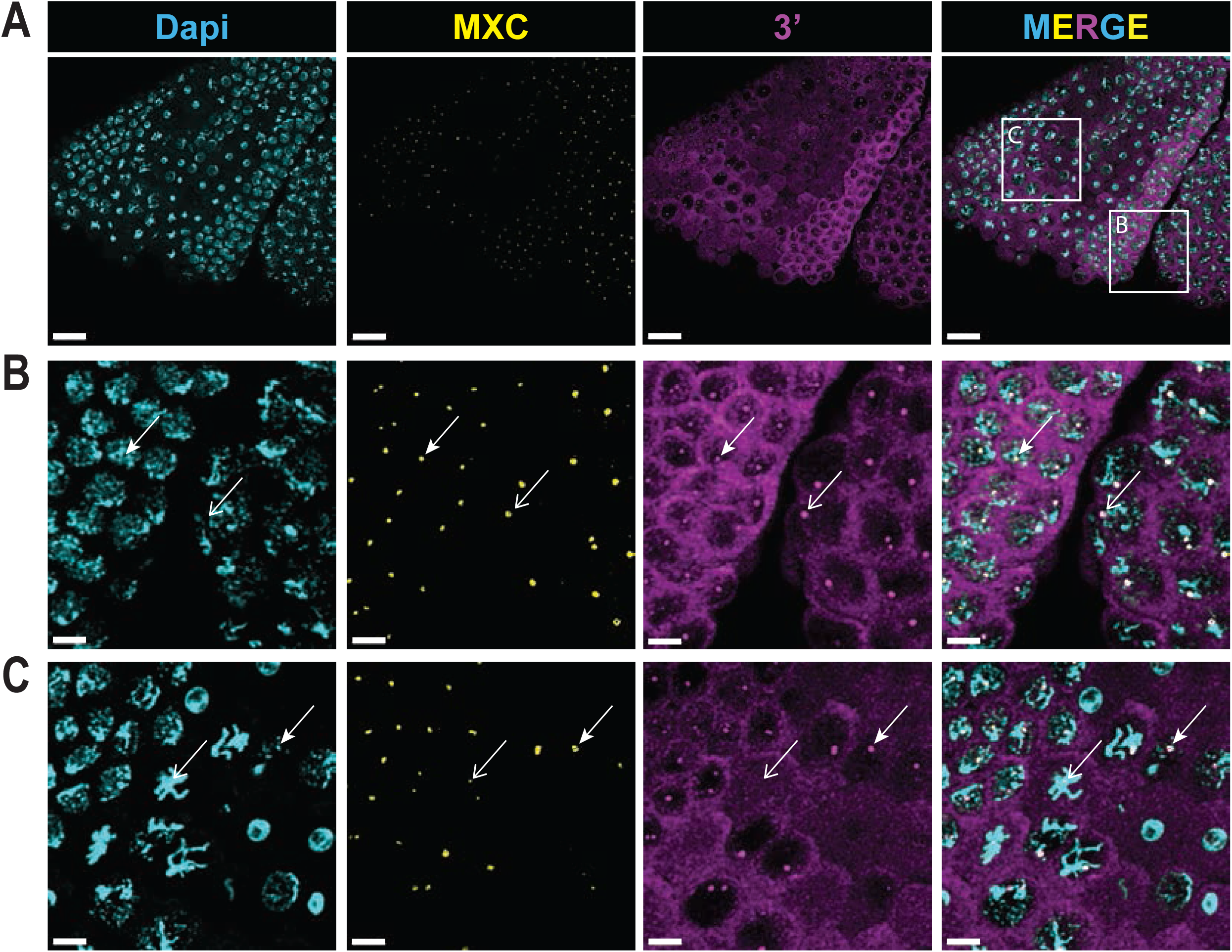
: **A)** Micrograph of the anterior portion of a gastrulating embryo stained for DAPI, Mxc, and the 3’ probe set. Notice the different mitotic domains highlighting G2, S, and M phases of the cell cycle in a single plane. Scale bars are 15 μM. **B)** Zoomed in region of interest “B” from the merge panel in A. The closed arrow points to an S-phase nucleus of cycle 15 and the open arrow highlights a G2 nucleus of cycle 14. Notice we can detect nascent 3’ transcripts in both nuclei demonstrating the continued transcription of the genes in both stages of the cell cycle at this stage in development. Scale bars are 4 μM. **C)** Zoomed in region of interest “C” from the merge panel in A. The closed arrow points to a nucleus in prophase demonstrating continued transcription as the cells enter mitosis. The open arrow points to a small Mxc focus on a metaphase chromosome, which lacks enriched FISH signal suggesting that transcription has been aborted by this point in mitosis. Scale bars are 4 μM.

Supplemental Videos 1: **A)** Time lapse of a GFP-MXC and mCherry-RPB1 expressing embryo through nuclear cycles 12-14. Same embryo as the micrographs from Figure 1B. GFP-MXC is cyan and mCherry-RPB1 is yellow. **B)** Same embryo as in A but only showing GFP-MXC. **C)** Same embryo as in A but only showing mCherry-RPB1.

Supplemental Videos 2: **A)** Time lapse of a GFP-MXC and mCherry-RPB1 expressing embryo as it enters gastrulation. Showing cells going from G2 through mitosis and into the next S-phase. Same embryo as Figure 1C. GFP-MXC is cyan and mCherry-RPB1 is yellow. **B)** Same embryo as in A but only showing GFP-MXC. **C)** Same embryo as in A but only showing mCherry-RPB1.

Supplemental Videos 3: **A)** Time lapse of a GFP-SPT6 and MXC-mScarlet expressing embryo through nuclear cycles 12-14. Same embryo as Figure 1D. GFP-SPT6 is cyan and MXC-mScarlet is yellow. **B)** Same embryo as in A but only showing GFP-SPT6. **C)** Same embryo as in A but only showing MXC-mScarlet.

Supplemental Videos 4: **A)** Time lapse of a GFP-SPT6 and mCherry-RPB1 expressing embryo through nuclear cycles 13-14. Same embryo as Figure 1E first panel. GFP-SPT6 is cyan and mCherry-RPB1 is yellow. **B)** Same embryo as in A but only showing GFP-SPT6. **C)** Same embryo as in A but only showing MXC-mScarlet.

Supplemental Videos 5: **A)** Time lapse of a GFP-SPT6 and mCherry-RPB1 expressing embryo through nuclear cycle 14 as the embryo enters gastrulation. Same embryo as Figure 1E second and third panels. GFP-SPT6 is cyan and mCherry-RPB1 is yellow. **B)** Same embryo as in A but only showing GFP-SPT6. **C)** Same embryo as in A but only showing MXC-mScarlet.

